# Rapid evolutionary turnover of mobile genetic elements drives microbial resistance to viruses

**DOI:** 10.1101/2021.03.26.437281

**Authors:** Fatima Aysha Hussain, Javier Dubert, Joseph Elsherbini, Mikayla Murphy, David VanInsberghe, Philip Arevalo, Kathryn Kauffman, Bruno Kotska Rodino-Janeiro, Martin Polz

**Author notes:** These authors contributed equally to this work (Fatima Aysha Hussain, Javier Dubert).

## Abstract

Although it is generally accepted that viruses (phages) drive bacterial evolution, how these dynamics play out in the wild remains poorly understood. Here we show that the arms race between phages and their hosts is mediated by large and highly diverse mobile genetic elements. These phage-defense elements display exceedingly fast evolutionary turnover, resulting in differential phage susceptibility among clonal bacterial strains while phage receptors remain invariant. Protection afforded by multiple elements is cumulative, and a single bacterial genome can harbor as many as 18 putative phage-defense elements. Overall, elements account for 90% of the flexible genome amongst closely related strains. The rapid turnover of these elements demonstrates that phage resistance is unlinked from other genomic features and that resistance to phage therapy might be as easily acquired as antibiotic resistance.

## Main Text

Bacterial viruses (phages) are ubiquitous across Earth’s biosphere and control microbial populations through predatory interactions (*1–3*). Because successful killing depends on molecular interaction with the host, phages display higher specificity than most other microbial predators—this being an important reason for renewed interest in the clinical use of phages as alternatives to antibiotics (*4*). Like antibiotics, however, phage killing exerts strong selection for resistance in bacterial hosts (*5*). Consequently, how bacteria naturally acquire resistance has important implications for understanding microbial community dynamics as well as the long-term success of phage therapy. Laboratory co-evolution studies have consistently identified phage-receptor mutations as key drivers of resistance (*6–8*), with genetic analyses suggesting secondary contributions by restriction-modification (RM) (*9–11*) and abortive infection (Abi) systems (*12, 13*). However, because phages target important surface structures such as the lipopolysaccharide (LPS) and membrane proteins as receptors, it is questionable whether mutations in receptors represent primary adaptive strategies in complex microbial communities since such modifications frequently incur significant fitness costs (*6*). Indeed, many additional defense mechanisms have recently been discovered, including clustered regularly interspaced short palindromic repeats (CRISPR) systems (*14, 15*), and several other yet-to-be mechanistically characterized defense mechanisms (*16, 17*). Importantly, genes encoding both receptors and defense systems have been shown to occur frequently in variable genomic islands (*6*, *18, 19*), or associated with mobile genetic elements (*20–22*). However, neither the predominant mechanisms of resistance to phages nor the dynamics of resistance gain and loss are well understood for microbes in the wild, limiting our knowledge of how phage predation structures diversity and drives evolution of microbial populations.

Here we combine population genomic and molecular genetic approaches to determine how phage resistance arises in microbial populations evolving in nature. We recently created the largest genomically-resolved phage-host cross-infection network, using sympatric environmental isolates, allowing us to ask at what genetic divergence and by what genetic mechanisms host resistance to phages arises in the wild (*23*, *24*). This system—the Nahant Collection—was established in the context of a 93-consecutive-day coastal-ocean time series and comprises over 1,300 strains of marine *Vibrio* ranging in relatedness from near-clonal to species-level divergence (*25*). *Vibrio* hosts isolated on three different days were used as “bait” to isolate 248 co-occurring lytic viruses by quantitative plaque assays, and each of 245 plaque positive hosts were subsequently challenged with all phage isolates to establish an all-by-all cross-infection matrix (Fig. S1A). Phages tend to be highly specific, as indicated by the general sparsity of the matrix. Additionally, even the most closely related hosts, differentiated by a few single nucleotide polymorphisms (SNPs) across the genome, are preyed upon by different phages (Fig. S1B). Furthermore, this extends to broader host-range phages that, although capable of infecting multiple hosts, are typically limited to a single strain within a host clade (Fig. S1C).

Because the observation that nearly clonal bacterial strains are subject to differential predation suggests extremely rapid evolution of phage resistance, we sought to identify the responsible mechanism in an exemplary set of 19, nearly clonal isolates of *Vibrio lentus.* These strains share identical nucleotide sequences in 52 ribosomal proteins (Fig. 1A, left) yet form two groups of 4 and 15 strains subject to differential infection by two groups of siphovirus phages consisting of 4 and 18 isolates (“purple” and “orange” in Fig. 1A, Fig. S2A-D). While the phage groups are so divergent that their genomes cannot be aligned (Fig. 1A, top; Fig. S2EF), the two host groups are so recently diverged that they differ only by 14 SNPs across their entire core genome (Fig. S3, Table S1). Re-assaying all pairwise interactions between the phages and hosts in this set over a wide range of phage-to-host ratios (multiplicities of infections, MOIs) revealed that representatives of both phage groups can also attach to strains originally scored as non-hosts and kill them at high concentrations, albeit without production of viable phage progeny (Fig. S4). This effect, termed “lysis from without” (*26*), along with observed adsorption of all phages to all hosts (Fig. S5), led us to hypothesize that all phages in these two groups can recognize receptors on all 19 bacterial strains and that differential resistance among these strains is mediated by intracellular mechanisms rather than by receptor modification.

**Fig. 1.**
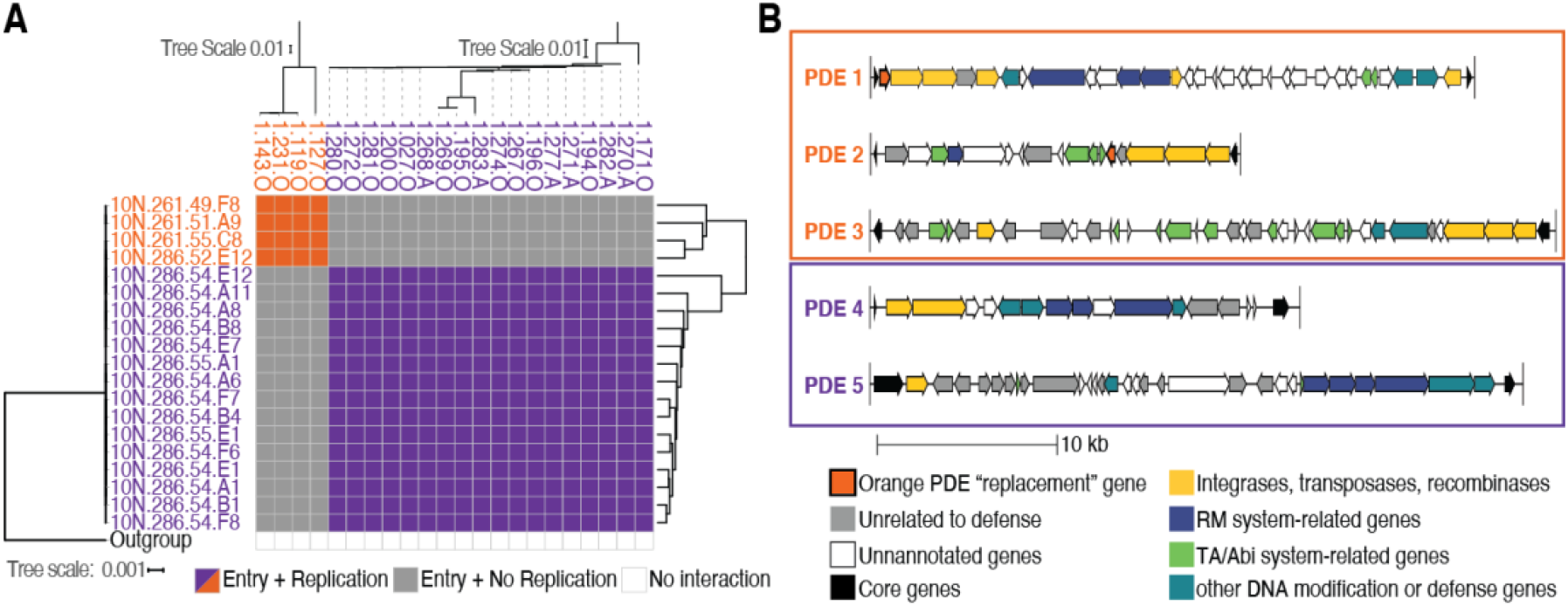
Near-clonal strains of *Vibrio lentus* differ in sensitivity to phage predation and differ in the carriage of mobile genetic elements encoding for phage-defense genes. **(A)** Phage host-range matrix with rows representing bacterial strains and columns representing phages. **(A-left)** Phylogenetic tree of 52 concatenated ribosomal protein sequences, **(A-top)** whole genome tree of viruses, **(A-right)** hierarchical clustering of whole genome alignments of bacterial hosts. **(B)** Gene diagrams of mobile genetic elements specific to the two host groups (as indicated by orange or purple outlines).

Supporting our hypothesis, we find that while the two phage types use different receptors, all bacterial genomes encode both sets of receptors. Transposon mutagenesis data suggests that the most likely candidate for the “orange” phage receptor is the Type II secretion system pseudopilus, while the “purple” phages likely use LPS as a primary receptor and a sodium transporter as a secondary receptor (Table S2). Despite the two phage groups using distinct receptors, every identified gene involved in phage entry is identical in nucleotide sequence across the two host types, and therefore part of their core genome. Additionally, SNPs identified upon selecting for laboratory evolved resistance in the host strains following phage exposure corroborated these results: SNPs in the same receptor loci were identified in the spontaneously resistant strains, verifying the receptor identification, and suggesting that the mode of SNP-based resistance evolution observed in the lab is not always representative of that in the wild (Table S3). All of this evidence supports our hypothesis that receptors do not drive phage specificity in these strains in the environment, leading us to explore potential intracellular mechanisms of host resistance by a combination of comparative genomics and molecular genetics.

Clustering the pangenome of *Vibrio lentus* isolates using high-quality genomes generated by hybrid assemblies of long and short reads (Fig. 1A, right) reveals that each host group harbors a set of putative phage-defense genes and that all of these genes are housed on large, genomically-integrated mobile genetic elements rather than merely being clustered in variable genomic regions as previously suggested for other defense genes (*19*). Three and two defined genomic regions specific to the “orange” and “purple” phage-susceptible hosts could be identified, respectively (Fig. 1B). These regions appear to be mobile genetic elements, which are likely transferred via site-specific recombination, because each element contains a defined insertion site on the host chromosome, potentially allowing for circularization, and all elements contain integrases and transposases on their periphery (Fig. 1B, Data S1). Similar elements can be found in very distantly related strains, suggesting their common horizontal acquisition and loss (Fig. S6). Because each element carries at least one unique known phage defense system, we refer to them as phage-defense elements (PDEs). PDE1, PDE4, and PDE5 have Type 1 RM systems, and PDE2 and PDE3 have putative toxin-antitoxin (TA) systems, some of which have been shown to act as Abi systems, killing the host upon phage infection (*27*). The PDEs range in size from approximately 10 to 60 kbp, and aside from defense systems and mobile element proteins, most remaining genes on the elements are unannotated. A striking feature of the PDEs is that their insertion does not appear to disrupt host functions. For example, PDE1 inserts into “orange” hosts’ 5’-deoxynucleotidase nucleosidase (Yfbr) gene, thereby truncating it, but encodes its own distinct copy of the same gene with a divergent amino acid sequence. Similarly, although PDE2 disrupts a thiol peroxidase gene upon insertion, it encodes for a second peroxidase copy in the middle of the element (Fig. 1B, Data S1).

That multiple putative PDEs cluster with phage resistance phenotypes poses the question: To what extent each of these elements contributes to the observed resistance? All three PDE-encoded RM systems appear active, as methylome data show that the distinct sequence motifs corresponding to each RM system are methylated only within “orange” and “purple” phage-host sets—that is, only in the bacterial genomes in which the RMs occur, and in the phages that can kill those hosts (Table S4). To further characterize the contribution to resistance of a full set of PDEs in the host populations, we systematically knocked out the putative defense portions of the three PDEs hypothesized to drive resistance of a representative “orange” host strain (10N.261.55.C8) to “purple” phages (Fig. S7A), and then challenged the knockouts with a representative “purple” phage (1.281.O) (Fig. 2A, Fig. S7B). This genetic analysis supports that in addition to the RM-encoding PDE specific to the “orange” host population, both PDEs encoding putative Abi systems are also active and contribute to phage resistance in a complex, cumulative manner. The complexity of the interaction among all PDEs leading to full resistance is illustrated by deleting them in all possible combinations. Knocking out RM-containing PDE1 alone increased killing tenfold (from 10^−1^ to 10^−2^ phage dilution), but still did not yield viable phage progeny (Fig. 2B). This phenotype is consistent with the expected host-killing of Abi systems (*28*) and led us to hypothesize that there may be a multi-level defense structure involving the remaining two PDEs as well. Knocking out Abi-system containing PDE2 and PDE3 independently resulted in no change from the wild type (WT) phenotype. However, knocking out PDE2 and PDE3 together allowed for both killing and propagation of the phage at a wider range of high MOIs (≤10^−5^ phage dilution). This phenotype is typical of an RM system-based defense strategy which is inherently imperfect (*29*)—at high MOIs, we expect higher numbers of co-infecting phages and thus an increase in the probability that an inadvertent phage-DNA methylation will occur at the target motif and allow a phage genome to escape restriction and replicate successfully. Knocking out PDE1 and PDE2 together and PDE1 and PDE3 together resulted in the same tenfold increase in killing observed when deleting PDE1 alone, indicating PDE2 and PDE3 provide a certain level of redundancy in protection. Finally, knocking out PDE1 and PDE3 together yields killing at much lower MOIs (10^−5^ phage dilution), suggesting PDE3 is a stronger resistance element than PDE2. However, the observed killing still did not consistently yield viable phage propagation (Fig. 2B). Finally, knocking out all three PDEs simultaneously resulted in the “orange” strain becoming just as susceptible to the “purple” phage as “purple” WT host strains. Therefore, we conclude all three elements are needed for full, WT-level defense.

**Fig. 2.**
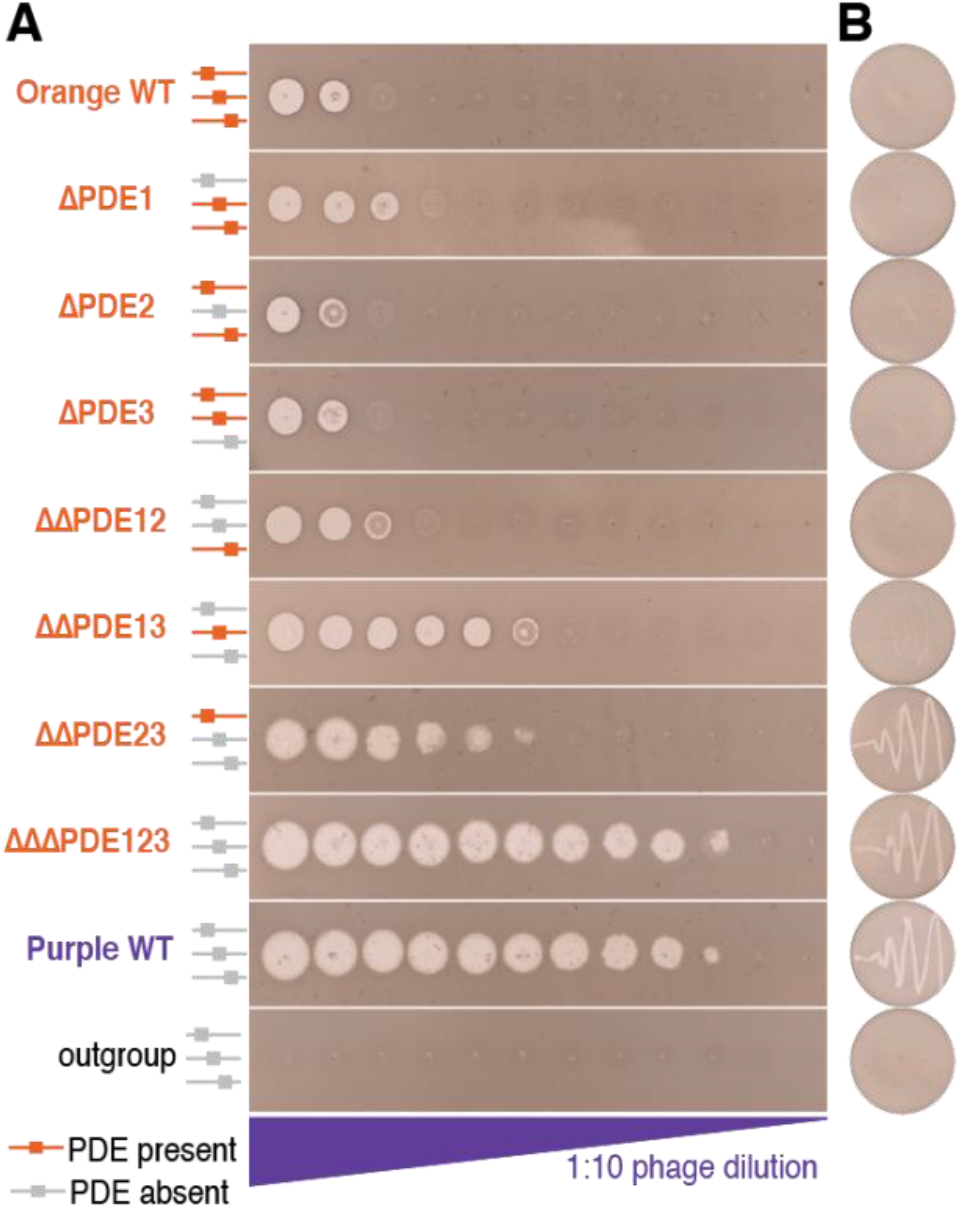
Changes in susceptibility to phage killing observed for phage-defense element (PDE) markerless deletions. (**A**) Lawns of bacterial hosts with drop spots of a 1:10 dilution series of “purple” phage (1.281.O). Cartoons on left indicate the presence or absence of different PDEs in each strain. From top to bottom: “orange” wild type host (10N.261.55.C8), ΔPDE1, ΔPDE2, ΔPDE3, ΔΔPDE12, ΔΔPDE13, ΔΔPDE23, ΔΔΔPDE123, “purple” wild type host (10N.286.54.F7, positive control), outgroup (10N.261.49.C11, negative control). (**B**) Re-streak test for propagation of phage progeny from drop spot clearings. Only infections of ΔΔPDE23, ΔΔΔPDE123, and “purple” wild type hosts produce viable phages, indicated by secondary clearing on the re-streak plates.

Expanding our genomic comparisons to additional closely related isolates in our collection, reveals that PDEs comprise the vast majority of the flexible gene content. Furthermore, PDEs account for a large fraction of the unannotated genes therein, thereby addressing the general open question of the function of the pan-genome (*30*). In addition to the 19 clones, we included 4 strains in our collection that are closely related but exhibit alternative phage sensitivity profiles. Using a k-mer-based approach to conduct all-by-all pair-wise genome comparisons (*31*), we identified a total of 30 unique putative PDEs, totaling 862,000 bp in length, shared by different subsets of the 23 strains analyzed (Fig. S8). The number of PDEs ranges between 10 and 18 in each strain and collectively the PDEs account for >90% of the flexible genomic regions, which can hence be given a tentative annotation (Fig. 3). Even if only known defense genes but not entire PDEs are considered, these account for 20% of the flexible gene content (Fig. 3). A similar range of 12-21% of flexible gene content is observed when we assayed the fraction of known defense genes in diverse other *Vibrio* species in our collection (Fig. S9). Importantly, because this measure only comprises the defense genes but not the entire PDEs, this suggests that a major portion, and perhaps the majority, of the pan-genome across these species is involved in phage defense and suggests a path forward for annotating this enigmatic genetic repertoire.

**Fig. 3.**
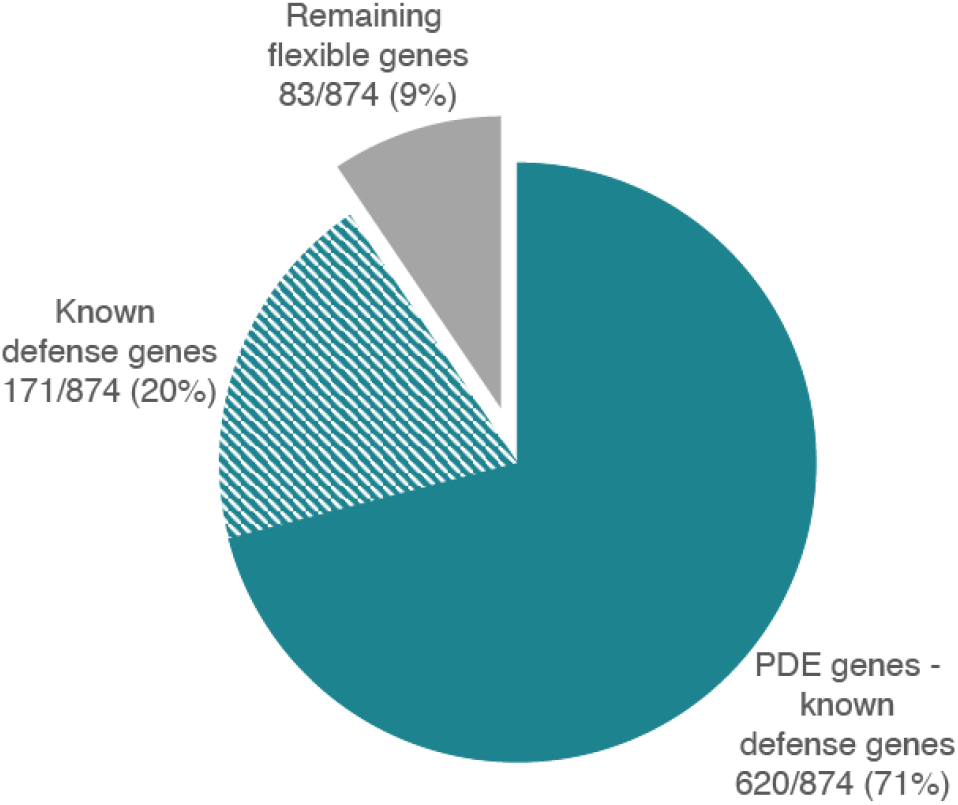
Fraction of the bacterial flexible genome attributed to phage defense. Amongst the 23 clones, an all-by-all genomic comparison shows 91% of flexible regions greater than 5kbp are putative PDEs. Only 20% of the PDEs match known defense genes while the remining are other PDE-specific genes, many of which are unannotated (71%).

Defense being entirely relegated to PDEs confirms theoretical considerations that resistance genes should be mobile because the cost of resistance limits their utility under changing predation pressures (*22*). Our findings demonstrate that the rate of turnover can be surprisingly fast, with only 14 SNPs accumulating across the entire genome per 5 PDE transfer events (gains or losses). This finding is likely more general as other *Vibrio* species in our collection, for which clonal isolates are available, also differ in their phage predation profiles, and even among the bacterial pathogens *Listeria* and *Salmonella* that follow a primarily non-recombinogenic mode of evolution, we observe similarly rapid turnover of putative PDEs (Fig. S10, Fig. S11). Thus, comparative genomics of near clonal isolates combined with phage host-range data is a fruitful method to discover novel phage-defense mechanisms in an unbiased way. However, we emphasize that high quality genomes assembled using long reads are essential since in our experience PDEs assemble poorly when using short read data, in part due to AT richness and high density of variable repeat regions.

Because receptor genes are invariant across the two near clonal host groups challenges the notion that receptor variation is primarily responsible for resistance (*6*, *19*), we asked to what extent the receptors identified are variable across more diverse populations. Populations are defined here as gene flow clusters that also represent ecological units (*32*, *33*), and the recognition of population boundaries is key for interpretation of gene or allele frequency in light of selective forces. Surprisingly, looking across 107 *Vibrio* isolates, spanning 10 populations, all putative receptors are highly monomorphic on the population level and possibly under purifying selection. The two genes identified as putative receptors in *V. lentus* are identical or nearly so at nucleotide level within most of the diverse populations and thus well below the average diversity of core genes (Fig. S12A). This is corroborated by phylogenetic trees showing that all members of each population carry the same or highly similar gene variants, a pattern consistent with recent gene-specific selective sweeps, with the notable exception of pseudopilin gene of the Type II secretion system, which is more diverse in two of the populations (Fig. S12B). Finally, the LPS, which frequently serves as primary receptor for many phages also appears similar at the population level since the genes responsible for synthesis display population-specific presence/absence patterns suggesting the synthesis pathway is conserved (Fig. S12C). Thus, although putative receptors can reside in variable regions when more divergent genomes are compared (*19*), their evolution appears much more constrained when population structure is considered. This constraint may arise because in wild populations, these surface structures are optimized for ecological interactions, and indicates a key difference from the lab where receptor mutations frequently arise in phage-host co-cultures (*34*). This observed invariance thus suggests that other selective forces that compete with phage resistance play an important role in receptor evolution in the wild and is consistent with predictions that intracellular defenses should be important under such conditions (*35*). It is possible that receptor-mediated defenses may only be advantageous under extreme predation regimes, or under regimes of low effective diversity such as a clonal infection.

The rapid turnover of PDEs implies that phage resistance is essentially unlinked from other traits within bacterial populations. Low linkage means that in complex microbial communities, bacterial core genomes can be maintained over the long-term even in the face of phage predation, while flexible genome content involved in shielding against phages is highly dynamic. In particular, our results question whether phage predation can increase microbial population diversity at the strain level by virtue of Kill-the-Winner type dynamics, which postulates that fitter genotypes are prevented from outcompeting all others within a population since they are disproportionally affected by phage predation (*36*). Instead, such dynamics are likely limited to acting at the resolution of mobile genetic elements and flexible genes, with limited consequences for the longer-term dynamics of bacterial population core genome diversity. This means that other factors, aside from phage predation, must drive the diversity observed in wild microbial populations. Similarly, rapid transfer of PDEs implies resistance to phage therapy may be easily acquired and quickly spread through bacterial populations, just as the connection to mobile genetic elements (primarily plasmids) has led to an unanticipated rise in antibiotic resistance. Together, our findings suggest that phage resistance is an important, if not the most important, selective force determining clonal bacterial diversity, with phage-defense elements potentially explaining a very large portion of the previously enigmatic bacterial pangenome.

## Acknowledgments

We thank D. Newman, K. Costa, H. Wildschutte, and F. Le Roux for advice and guidance with mutagenesis experiments. We thank M. Cutler for experimental support. We thank S. W. Chisholm and L. Kelly for thoughtful comments. We thank O. X. Cordero, S. Kearney, and A. F. Takemura for valuable suggestions throughout.

## Author contributions

F.A.H. and M.P. conceived of the project, designed the study, and wrote the paper with contributions from all coauthors. F.A.H. performed the experiments with assistance from J.D. and M.M.. F.A.H., J.D., and B.R. designed the genetic knockouts and conducted manual annotations. J.D. made the knockout mutants. F.A.H. curated the data, conducted formal analyses, and prepared the figures with feedback from all coauthors. F.A.H., J.E., M.M., P.A., and D.V. wrote and/or optimized code for the bioinformatic pipelines. K.K. isolated and performed initial characterization of the lytic viruses. M.P. supervised the project and secured funding.

## Data and materials availability

New genomes used in this work have been deposited under the NCBI BioProject with accession number PRJNA328102. All data, code, and materials are available upon request.

## References and Notes

1. K. E. Wommack, R. R. Colwell, Virioplankton: Viruses in Aquatic Ecosystems. Microbiol. Mol. Biol. Rev. 64, 69–114 (2000).

2. M. Breitbart, F. Rohwer, Here a virus, there a virus, everywhere the same virus? Trends Microbiol. 13, 278–284 (2005).

3. H. G. Hampton, B. N. J. Watson, P. C. Fineran, The arms race between bacteria and their phage foes. Nature. 577, 327–336 (2020).

4. K. E. Kortright, B. K. Chan, J. L. Koff, P. E. Turner, Phage Therapy: A Renewed Approach to Combat Antibiotic-Resistant Bacteria. Cell Host Microbe. 25, 219–232 (2019).

5. S. J. Labrie, J. E. Samson, S. Moineau, Bacteriophage resistance mechanisms. Nat. Rev. Microbiol. 8, 317–327 (2010).

6. S. Avrani, O. Wurtzel, I. Sharon, R. Sorek, D. Lindell, Genomic island variability facilitates Prochlorococcus-virus coexistence. Nature. 474, 604–608 (2011).

7. J. R. Meyer, D. T. Dobias, J. S. Weitz, J. E. Barrick, R. T. Quick, R. E. Lenski, Repeatability and Contingency in the Evolution of a Key Innovation in Phage Lambda. Science. 335, 428–432 (2012).

8. B. j. m. Bohannan, R. e. Lenski, Linking genetic change to community evolution: insights from studies of bacteria and bacteriophage. Ecol. Lett. 3, 362–377 (2000).

9. W. Arber, D. Dussoix, Host specificity of DNA produced by Escherichia coli: I. Host controlled modification of bacteriophage λ. J. Mol. Biol. 5, 18–36 (1962).

10. G. G. Wilson, N. E. Murray, Restriction and Modification Systems. Annu. Rev. Genet. 25, 585–627 (1991).

11. G. Bertani, J. J. Weigle, Host Controlled Variation in Bacterial Viruses. J. Bacteriol. 65, 113–121 (1953).

12. I. J. Molineux, Host-parasite interactions: recent developments in the genetics of abortive phage infections. New Biol. 3, 230–236 (1991).

13. L. Snyder, Phage-exclusion enzymes: a bonanza of biochemical and cell biology reagents? Mol. Microbiol. 15, 415–420 (1995).

14. R. Barrangou, C. Fremaux, H. Deveau, M. Richards, P. Boyaval, S. Moineau, D. A. Romero, P. Horvath, CRISPR Provides Acquired Resistance Against Viruses in Prokaryotes. Science. 315, 1709–1712 (2007).

15. A. F. Andersson, J. F. Banfield, Virus Population Dynamics and Acquired Virus Resistance in Natural Microbial Communities. Science. 320, 1047–1050 (2008).

16. S. Doron, S. Melamed, G. Ofir, A. Leavitt, A. Lopatina, M. Keren, G. Amitai, R. Sorek, Systematic discovery of antiphage defense systems in the microbial pangenome. Science, eaar4120 (2018).

17. R. P. Novick, G. E. Christie, J. R. Penadés, The phage-related chromosomal islands of Gram positive bacteria. Nat. Rev. Microbiol. 8, 541–551 (2010).

18. K. S. Makarova, Y. I. Wolf, S. Snir, E. V. Koonin, Defense Islands in Bacterial and Archaeal Genomes and Prediction of Novel Defense Systems. J. Bacteriol. 193, 6039–6056 (2011).

19. F. Rodriguez-Valera, A.-B. Martin-Cuadrado, B. Rodriguez-Brito, L. Pašić, T. F. Thingstad, F. Rohwer, A. Mira, Explaining microbial population genomics through phage predation. Nat. Rev. Microbiol. 7, 828–836 (2009).

20. N. D. McDonald, A. Regmi, D. P. Morreale, J. D. Borowski, E. F. Boyd, CRISPR-Cas systems are present predominantly on mobile genetic elements in Vibrio species. BMC Genomics. 20, 105 (2019).

21. A. C. McKitterick, K. D. Seed, Anti-phage islands force their target phage to directly mediate island excision and spread. Nat. Commun. 9, 2348 (2018).

22. E. V. Koonin, K. S. Makarova, Y. I. Wolf, M. Krupovic, Evolutionary entanglement of mobile genetic elements and host defence systems: guns for hire. Nat. Rev. Genet., 1–13 (2019).

23. K. M. Kauffman, F. A. Hussain, J. Yang, P. Arevalo, J. M. Brown, W. K. Chang, D. VanInsberghe, J. Elsherbini, R. S. Sharma, M. B. Cutler, L. Kelly, M. F. Polz, A major lineage of non-tailed dsDNA viruses as unrecognized killers of marine bacteria. Nature. 554, 118 (2018).

24. K. M. Kauffman, J. M. Brown, R. S. Sharma, D. VanInsberghe, J. Elsherbini, M. Polz, L. Kelly, Viruses of the Nahant Collection, characterization of 251 marine Vibrionaceae viruses. Sci. Data. 5, 180114 (2018).

25. A. M. Martin-Platero, B. Cleary, K. Kauffman, S. P. Preheim, D. J. McGillicuddy, E. J. Alm, M. F. Polz, High resolution time series reveals cohesive but short-lived communities in coastal plankton. Nat. Commun. 9, 266 (2018).

26. M. Delbrück, The Growth of Bacteriophage and Lysis of the Host. J. Gen. Physiol. 23, 643–660 (1940).

27. R. L. Dy, C. Richter, G. P. C. Salmond, P. C. Fineran, Remarkable Mechanisms in Microbes to Resist Phage Infections. Annu. Rev. Virol. 1, 307–331 (2014).

28. G. Lindahl, G. Sironi, H. Bialy, R. Calendar, Bacteriophage Lambda; Abortive Infection of Bacteria Lysogenic for Phage P2. Proc. Natl. Acad. Sci. 66, 587–594 (1970).

29. D. H. Kruger, T. A. Bickle, Bacteriophage survival: multiple mechanisms for avoiding the deoxyribonucleic acid restriction systems of their hosts. Microbiol. Rev. 47, 345–360 (1983).

30. E. P. C. Rocha, Neutral Theory, Microbial Practice: Challenges in Bacterial Population Genetics. Mol. Biol. Evol. 35, 1338–1347 (2018).

31. Materials, methods, and supplemental text are available as supplementary materials on Science Online.

32. P. Arevalo, D. VanInsberghe, J. Elsherbini, J. Gore, M. F. Polz, A Reverse Ecology Approach Based on a Biological Definition of Microbial Populations. Cell. 178, 820–834.e14 (2019).

33. B. J. Shapiro, J. Friedman, O. X. Cordero, S. P. Preheim, S. C. Timberlake, G. Szabó, M. F. Polz, E. J. Alm, Population Genomics of Early Events in the Ecological Differentiation of Bacteria. Science. 336, 48–51 (2012).

34. E. R. Westra, S. van Houte, S. Oyesiku-Blakemore, B. Makin, J. M. Broniewski, A. Best, J. Bondy-Denomy, A. Davidson, M. Boots, A. Buckling, Parasite Exposure Drives Selective Evolution of Constitutive versus Inducible Defense. Curr. Biol. 25, 1043–1049 (2015).

35. S. Zborowsky, D. Lindell, Resistance in marine cyanobacteria differs against specialist and generalist cyanophages. Proc. Natl. Acad. Sci., 201906897 (2019).

36. C. Winter, T. Bouvier, M. G. Weinbauer, T. F. Thingstad, Trade-Offs between Competition and Defense Specialists among Unicellular Planktonic Organisms: the “Killing the Winner” Hypothesis Revisited. Microbiol. Mol. Biol. Rev. 74, 42–57 (2010).

37. D. E. Hunt, L. A. David, D. Gevers, S. P. Preheim, E. J. Alm, M. F. Polz, Resource Partitioning and Sympatric Differentiation Among Closely Related Bacterioplankton. Science. 320, 1081–1085 (2008).

38. S. G. John, C. B. Mendez, L. Deng, B. Poulos, A. K. M. Kauffman, S. Kern, J. Brum, M. F. Polz, E. A. Boyle, M. B. Sullivan, A simple and efficient method for concentration of ocean viruses by chemical flocculation. Environ. Microbiol. Rep. 3, 195–202 (2011).

39. K. M. Kauffman, M. F. Polz, Streamlining standard bacteriophage methods for higher throughput. MethodsX. 5, 159–172 (2018).

40. A. K. M. Kauffman, thesis, Massachusetts Institute of Technology (2014).

41. S. R. Eddy, Accelerated Profile HMM Searches. PLoS Comput. Biol. 7 (2011), doi:10.1371/journal.pcbi.1002195.

42. K. Katoh, D. M. Standley, MAFFT Multiple Sequence Alignment Software Version 7: Improvements in Performance and Usability. Mol. Biol. Evol. 30, 772–780 (2013).

43. A. Stamatakis, RAxML version 8: a tool for phylogenetic analysis and post-analysis of large phylogenies. Bioinformatics. 30, 1312–1313 (2014).

44. N.-F. Alikhan, N. K. Petty, N. L. Ben Zakour, S. A. Beatson, BLAST Ring Image Generator (BRIG): simple prokaryote genome comparisons. BMC Genomics. 12, 402 (2011).

45. L. A. Kelley, S. Mezulis, C. M. Yates, M. N. Wass, M. J. E. Sternberg, The Phyre2 web portal for protein modeling, prediction and analysis. Nat. Protoc. 10, 845–858 (2015).

46. J. Huerta-Cepas, K. Forslund, L. P. Coelho, D. Szklarczyk, L. J. Jensen, C. von Mering, P. Bork, Fast Genome-Wide Functional Annotation through Orthology Assignment by eggNOG-Mapper. Mol. Biol. Evol. 34, 2115–2122 (2017).

47. J. Huerta-Cepas, D. Szklarczyk, D. Heller, A. Hernández-Plaza, S. K. Forslund, H. Cook, D. R. Mende, I. Letunic, T. Rattei, L. J. Jensen, C. von Mering, P. Bork, eggNOG 5.0: a hierarchical, functionally and phylogenetically annotated orthology resource based on 5090 organisms and 2502 viruses. Nucleic Acids Res. 47, D309–D314 (2019).

48. L. Guy, J. Roat Kultima, S. G. E. Andersson, genoPlotR: comparative gene and genome visualization in R. Bioinformatics. 26, 2334–2335 (2010).

49. M. Steinegger, J. Söding, MMseqs2 enables sensitive protein sequence searching for the analysis of massive data sets. Nat. Biotechnol. 35, 1026–1028 (2017).

50. J. P. Meier-Kolthoff, M. Göker, VICTOR: genome-based phylogeny and classification of prokaryotic viruses. Bioinformatics. 33, 3396–3404 (2017).

51. M. Baym, S. Kryazhimskiy, T. D. Lieberman, H. Chung, M. M. Desai, R. Kishony, Inexpensive Multiplexed Library Preparation for Megabase-Sized Genomes. PLOS ONE. 10, e0128036 (2015).

52. https://github.com/rrwick/Filtlong.

53. M. Kolmogorov, J. Yuan, Y. Lin, P. A. Pevzner, Assembly of long, error-prone reads using repeat graphs. Nat. Biotechnol. 37, 540–546 (2019).

54. R. R. Wick, M. B. Schultz, J. Zobel, K. E. Holt, Bandage: interactive visualization of de novo genome assemblies. Bioinformatics. 31, 3350–3352 (2015).

55. http://www.bioinformatics.babraham.ac.uk/projects/trim_galore/.

56. R. R. Wick, L. M. Judd, C. L. Gorrie, K. E. Holt, Unicycler: Resolving bacterial genome assemblies from short and long sequencing reads. PLOS Comput. Biol. 13, e1005595 (2017).

57. D. Hyatt, G.-L. Chen, P. F. LoCascio, M. L. Land, F. W. Larimer, L. J. Hauser, Prodigal: prokaryotic gene recognition and translation initiation site identification. BMC Bioinformatics. 11, 119 (2010).

58. P. Jones, D. Binns, H.-Y. Chang, M. Fraser, W. Li, C. McAnulla, H. McWilliam, J. Maslen, A. Mitchell, G. Nuka, S. Pesseat, A. F. Quinn, A. Sangrador-Vegas, M. Scheremetjew, S.-Y. Yong, R. Lopez, S. Hunter, InterProScan 5: genome-scale protein function classification. Bioinformatics. 30, 1236–1240 (2014).

59. N. Yutin, P. Puigbò, E. V. Koonin, Y. I. Wolf, Phylogenomics of Prokaryotic Ribosomal Proteins. PLoS ONE. 7 (2012), doi:10.1371/journal.pone.0036972.

60. T. J. Treangen, B. D. Ondov, S. Koren, A. M. Phillippy, The Harvest suite for rapid core genome alignment and visualization of thousands of intraspecific microbial genomes. Genome Biol. 15, 524 (2014).

61. L. Ferrières, G. Hémery, T. Nham, A.-M. Guérout, D. Mazel, C. Beloin, J.-M. Ghigo, Silent Mischief: Bacteriophage Mu Insertions Contaminate Products of Escherichia coli Random Mutagenesis Performed Using Suicidal Transposon Delivery Plasmids Mobilized by Broad-Host-Range RP4 Conjugative Machinery. J. Bacteriol. 192, 6418–6427 (2010).

62. S. L. Chiang, E. J. Rubin, Construction of a mariner-based transposon for epitope-tagging and genomic targeting. Gene. 296, 179–185 (2002).

63. F. Le Roux, J. Binesse, D. Saulnier, D. Mazel, Construction of a Vibrio splendidus Mutant Lacking the Metalloprotease Gene vsm by Use of a Novel Counterselectable Suicide Vector. Appl. Environ. Microbiol. 73, 777–784 (2007).

64. S. Das, J. C. Noe, S. Paik, T. Kitten, An improved arbitrary primed PCR method for rapid characterization of transposon insertion sites. J. Microbiol. Methods. 63, 89–94 (2005).

65. F. M. Lauro, K. Tran, A. Vezzi, N. Vitulo, G. Valle, D. H. Bartlett, Large-Scale Transposon Mutagenesis of Photobacterium profundum SS9 Reveals New Genetic Loci Important for Growth at Low Temperature and High Pressure. J. Bacteriol. 190, 1699–1709 (2008).

66. Y. Jiao, A. Kappler, L. R. Croal, D. K. Newman, Isolation and Characterization of a Genetically Tractable Photoautotrophic Fe(II)-Oxidizing Bacterium, Rhodopseudomonas palustris Strain TIE-1. Appl. Environ. Microbiol. 71, 4487–4496 (2005).

67. C. Camacho, G. Coulouris, V. Avagyan, N. Ma, J. Papadopoulos, K. Bealer, T. L. Madden, BLAST+: architecture and applications. BMC Bioinformatics. 10, 421 (2009).

68. S. V. Angiuoli, S. L. Salzberg, Mugsy: fast multiple alignment of closely related whole genomes. Bioinforma. Oxf. Engl. 27, 334–342 (2011).

69. R Core Team, R: A Language and Environment for Statistical Computing (R Foundation for Statistical Computing, Vienna, Austria, 2019; https://www.R-project.org).

70. L. Zimmermann, A. Stephens, S.-Z. Nam, D. Rau, J. Kübler, M. Lozajic, F. Gabler, J. Söding, A. N. Lupas, V. Alva, A Completely Reimplemented MPI Bioinformatics Toolkit with a New HHpred Server at its Core. J. Mol. Biol. 430, 2237–2243 (2018).

71. S. F. Altschul, T. L. Madden, A. A. Schaffer, J. Zhang, Z. Zhang, W. Miller, D. J. Lipman, Gapped BLAST and PSI-BLAST: a new generation of protein database search programs. Nucleic Acids Res. 25, 3389–3402 (1997).

72. I. A. Murray, T. A. Clark, R. D. Morgan, M. Boitano, B. P. Anton, K. Luong, A. Fomenkov, S. W. Turner, J. Korlach, R. J. Roberts, The methylomes of six bacteria. Nucleic Acids Res., gks891 (2012).

73. R. J. Roberts, T. Vincze, J. Posfai, D. Macelis, REBASE--a database for DNA restriction and modification: enzymes, genes and genomes. Nucleic Acids Res. 43, D298–299 (2015).

74. J. Söding, A. Biegert, A. N. Lupas, The HHpred interactive server for protein homology detection and structure prediction. Nucleic Acids Res. 33, W244–W248 (2005).

75. M.-E. Val, O. Skovgaard, M. Ducos-Galand, M. J. Bland, D. Mazel, Genome Engineering in Vibrio cholerae: A Feasible Approach to Address Biological Issues. PLOS Genet. 8, e1002472 (2012).

76. G. Marçais, C. Kingsford, A fast, lock-free approach for efficient parallel counting of occurrences of k-mers. Bioinformatics. 27, 764–770 (2011).

77. B. Langmead, S. L. Salzberg, Fast gapped-read alignment with Bowtie 2. Nat. Methods. 9, 357–359 (2012).

78. B. D. Ondov, T. J. Treangen, P. Melsted, A. B. Mallonee, N. H. Bergman, S. Koren, A. M. Phillippy, Mash: fast genome and metagenome distance estimation using MinHash. Genome Biol. 17, 132 (2016).

79. M. Bastian, S. Heymann, M. Jacomy, in Third International AAAI Conference on Weblogs and Social Media (2009; https://www.aaai.org/ocs/index.php/ICWSM/09/paper/view/154).

80. R. C. Edgar, MUSCLE: multiple sequence alignment with high accuracy and high throughput. Nucleic Acids Res. 32, 1792–1797 (2004).

81. H. Wildschutte, S. P. Preheim, Y. Hernandez, M. F. Polz, O-antigen diversity and lateral transfer of the wbe region among Vibrio splendidus isolates. Environ. Microbiol. 12, 2977–2987 (2010).

