## Supplementary Materials for "Rapid evolutionary turnover of mobile genetic elements drives microbial resistance to viruses"

### Materials and Methods

Bacteria and phage isolation

Bacteria and phages were obtained in a previous study from coastal seawater collected from Canoe Cove, Nahant, MA, USA, on August 22 (ordinal day 222), September 18 (261), and October 13 (286), 2010 (*23*). *Vibrio* bacteria were isolated using a size fractionation approach, followed by plating on selective media, as described previously (*37*). Briefly, to capture bacteria associated with large particles and zoo- and phytoplankton, seawater was filtered through a 63 µm average pore size plankton net. To capture bacteria in smaller size fractions, including small particles and smaller zoo- and phytoplankton as well as bacteria occurring in the free-living fraction, water pre-filtered through a 63 µm net was serially passed through 5 µm, 1 µm, and 0.2 µm polycarbonate filters. To isolate vibrios from each fraction, material captured in the plankton net and on filters was resuspended in artificial seawater (ASW; Sea Salts from Sigma-Aldrich), and the suspensions passed through polyethersulfone 0.2 µm filters. These final filters were placed directly on agar plates of MTCBS (Difco Thiosulfate-Citrate-Bile-Sucrose Agar amended with 10 g/L of NaCl to final concentration of 2% w/v) to allow for selective growth of *Vibrio* colonies. Colonies were purified by serial passaging on agar plates of: first, TSB2 (Tryptic Soy Broth, 1.5% Difco Bacto Agar, amended with 15 g NaCl to 2% w/v); second, MTCBS, and third TSB2. Colonies were inoculated into 1 mL of Difco 2216 Marine Broth (2216MB) in 96-well 2 mL culture blocks and allowed to grow, shaking at room temperature, for 48 hours. Glycerol stocks for preservation at -80°C were prepared by combining 100 µL of culture with 100 µL of 50% glycerol (50% water) in 96-well microtiter plates. The naming of each strain reflects isolation location, day, and size fraction: 10N refers to the 2010 collection of samples from Nahant; 222, 261, 286 are the ordinal dates of the year; 54, 55, 56 are three replicates of the 63 µm fraction; 51, 52, 53, are three replicates of the 5 µm fraction; 48, 49, 50 are three replicates of the 1 µm fraction; and 45, 46, 47 are three replicates of the 0.2 µm or free-living fraction. The final portion of the name is the original storage well in a 96-well plate. Note that the "orange" isolates were, with one exception, collected on day 261 and distributed between the 1 µm, 5 µm, and 63 µm fraction, while all "purple" isolates were collected on day 286 from the 63 µm fraction. Therefore, the dynamics described in this work are likely occurring in particle-attached bacterial hosts in the ocean.

For phage isolation, 4 L of seawater was collected in triplicate on each day in the time series and separately filtered through a Sterivex 0.22 µm barrel filter into a sterile 4 L collection bottle using a peristaltic pump. Phages were directly concentrated from this filtrate using an iron flocculation and filtering method described previously (*38*). Briefly, iron (III) chloride, which is spiked into the sample, precipitates phages from the solution, and then the precipitates are collected onto 90 mm 0.2 µm polycarbonate filters using a glass cup-frit system. Precipitates are finally dissolved in 4 mL of oxalate solution to yield a quantitative concentration of 1,000x from the original 4 L. The final phage concentrate was stored at 4ºC in the dark until used to isolate specific viruses for different bacterial hosts.

*Vibrio* isolates were used as "bait" to obtain phages from the concentrates using direct plating in soft agar overlays. Plaques from the bait assay were archived frozen in 2216MB and glycerol and phages for use in the host range assay were subsequently randomly selected from archives for each host and purified by triple serial passage using tube-free agar overlays (*24*, *39*) on their hosts of isolation. Phages were amplified on their hosts of isolation using primary small-scale liquid cultures inoculated with plaque plugs from their final serial passage in agar overlays, followed by plating of primary lysates into agar overlays to achieve high-titer stocks. Top agar layers of lawns "at confluence" (saturated with plaques but not completely cleared) were harvested into 2216MB, centrifuged at 5,000 x *g* for 20 minutes, and filtered through Sterivex 0.22 µm barrel filters to generate the lysates used for the all-by-all host range cross test as well as for phage DNA extraction and sequencing (*24*), as well as methylation profiling (described below).

Phage host-range matrix

Phage host-range was determined in a previous study (*23*). Briefly, all *Vibrio* strains for which at least one phage was found (“plaque positive”) were used in the host-range assay and challenged with all phages purified as described above. Bacterial hosts were plated in agar overlays in large 150 mm plates and stamped with phage lysates arranged in triplicate in 96-well arrays using 96-spot blotters (BelArt, Bel-blotter 96-tip replicator, 378760002). Clearing seen in at least 2/3 replicates was scored as a positive kill (*40*). The concentration of each lysate in the original assay was not normalized to allow for higher throughput but the assay was repeated for select hosts at a range of concentrations (see methods for “varying phage concentrations” below).

To organize bacterial hosts in the matrix by phylogeny, concatenation of ribosomal proteins and *hsp60* sequences was used to construct a phylogenetic tree reflecting the relationship of the core genome (Fig. S1A). When genome sequences were available, we used HMMER (*41*) to find ribosomal proteins, and aligned the sequences with MAFFT (*42*). Amino acid sequences of *hsp60* proteins were also extracted from genomes via HMMER using pfam PF00118. The *hsp60* sequences were aligned using the mafft-fftnsi algorithm. When genomes were not available, *hsp60* sequences that were Sanger-sequenced were added to this alignment using the mafft-fftnsi algorithm with the –addfragments option. The *hsp60* alignment and the ribosomal protein alignment were concatenated and used to create the phylogenetic tree in Figure S1A using RAxML (options: –q, -m GTRGAMMAX) (*43*).

Phage characterization

Circular representations of the previously sequenced phage genomes (Fig. S2CD) were generated using BRIG (*44*) with publicly available NCBI GenBank files; annotations were made based on manual review of GenBank predictions and supplemented with Phyre2 (*45*) and EggNog-Mapper (*46*, *47*) annotation. Genome diagrams (Fig. S2EF) were generated using the GenoPlotR package in R (*48*) with predicted protein coding genes indicated as arrows colored to correspond to protein sequence clusters, as defined using default settings of MMseqs2 (*49*); and with “orange” (Fig. S2E) and “purple” (Fig. S2F) phages clustered and identified as two separate genus-level groups using the D6 amino acid OPTSIL clustering algorithm in the VICTOR classifier (*50*) with whole genome concatenated protein sequences.

Hybrid assemblies of bacterial genomes

Because we noticed that PDEs assemble poorly when genomes were sequenced with short read technology, we used Illumina short reads and Pacific Biosciences (PacBio) long reads in combination to assemble high quality (nearly closed) genomes. For short read sequencing, bacterial isolates were grown overnight from a single colony in 1.2 mL of 2216MB in deep-well blocks and processed in bulk. Genome libraries were prepared for sequencing using the Nextera DNA Library Preparation Kit (Illumina) with 1-2 ng input DNA per isolate, as previously described (*51*). Genomes were sequenced on 100 bp pared-end sequencing runs using Illumina HiSeq, with 50-60 samples multiplexed per lane. When available, Illumina HiSeq short reads from previous work (*23*) were used, otherwise new short read data were generated for this study. High quality bacterial genomic DNA for PacBio sequencing was prepared separately. A single colony was inoculated into a 250 mL Erlenmeyer flask with 50 mL of 2216MB and grown shaking at room temperature for 24 hours. The fresh culture was pelleted by centrifugation at 5,000 x *g* for 20 minutes and then immediately processed for DNA extraction when possible, or frozen at -20°C for short term storage. The Qiagen Genomic Tip 500x kit was used following the manufactures guidelines, and the final DNA was collected by spooling using a glass rod rather than centrifugation to avoid shearing. DNA was stored in 500 µL of elution buffer at 4°C for 24-48 hours to allow for full resuspension before sequencing at either the Yale Center for Genome Analysis (PacBio RS II, without multiplexing) or the BioMicroCenter at MIT (Sequel, with multiplexing).

A custom hybrid assembly pipeline was designed to process the data. Briefly, Pacbio reads were filtered at different length cutoffs using Filtlong (*52*) and then assembled using Flye (*53*) to create a set of reference genomes. The reference genomes were visualized using Bandage (*54*) and the best genome was selected, based on completion and coverage, to be used as a reference in the final assembly. For the final assembly, Illumina reads were trimmed using Trim-Galore (*55*), Pacbio reads were quality filtered with the trimmed Illumina reads using Filtlong, and both sets of processed reads, together with the best Flye assembly, were used as inputs for the Unicycler (*56*) assembler.

Bacterial genome annotations

Genomes were annotated using Prodigal 2.6 (*57*) for Open Reading Frame (ORF) prediction. Predicted ORFs were annotated using InterProScan5 (*58*) using the iprlookup, goterms, and pathways options. InterProScan5 matches against 13 databases by default, which are listed here: https://github.com/ebi-pf-team/interproscan/wiki/HowToRun#included-analyses. Two optional databases were included for this analysis: TMHMM for predicted transmembrane proteins and SignalP for predicted signal peptide cleavage sites.

Host relationships

Phylogenetic relationships among the 19 “orange” and “purple” clones were estimated by comparison of (i) concatenated alignments of ribosomal proteins and *hsp60* as described above in methods for, “Phage host-range matrix” (Fig. S1A), (ii) nucleotide sequences of 52 core ribosomal proteins (Fig. 1A, left) (*59*), and (iii) all shared genes (Fig. S3). For ribosomal protein comparisons, we searched for ribosomal proteins in the different genomes using HMMER (*41*), filtered the hits using custom python scripts, aligned the hits using MAFFT (*42*), concatenated the alignment using custom python scripts, and constructed the tree using RAxML (parameters raxmlHPC-PTHREADS -f a -x 26789416 -m GTRGAMMAX -p 218957 -# 100) (*43*). For estimation of whole genome relationships, we used the Parsnp  program (*60*) with the recombination flag (-x) to construct whole genome SNP trees. Then, HarvestTools was used to convert from a ggr format to a snp fasta file, and finally, IQ-Tree was used to optimize the final tree (Fig. S3). SNPs in the core genome were located using custom python scripts and verified by visualizing on Ginger, the Harvest graphic user interface. SNP details are given in Table S1.

Host range assays at varying phage concentrations

In order to determine the host-ranges of the specific phages used in this work at a higher resolution, we re-assayed a subset of hosts and phages of interest using a range of concentrations. Bacterial hosts were grown in 5 mL of 2216MB overnight from single colonies streaked on 1.5% Bacto Agar plates supplemented with 2216MB. Phage lysates were prepared as described above and diluted in 2216MB to form a ten-fold dilution series from $10^{0}$-$10^{-7}$. Five μL drop spots of each dilution were pipetted onto bacterial host lawns made using a tube-free agar overlay method (*39*) and incubated at room temperature for 24 hours before evaluating phage entry and efficiency of plating at the varying concentrations (Fig.1A). Plates with different killing were imaged using a flatbed scanner (Epson Perfection V800 Photo Scanner - Product No. B11B22320) and captured using VueScan Software by Hamrick (Fig. S4). Lysis from without was only observed when using this higher resolution assay, thus the authors recommend performing such an assay when evaluating phage host-range whenever possible.

Phage adsorption assay

In order to determine if all phages were attaching to all hosts, for “orange” phage 1.143.O and “purple” phage 1.281.O, we compared the number of free phages remaining in solution after exposure to “orange” host 10N.261.55.C8, “purple” host 10N.286.54.F7, an unrelated *Vibrio* (outgroup) control 10N.261.49.C11, and a no-host negative control (2216MB). Three different colonies of each bacterial strain were inoculated in 3 mL of 2216MB and grown shaking at 25°C for 4 hours. Bacterial concentration was estimated at optical density measured at 600 nm wavelength ($OD_{600}$), and each replicate was normalized to $OD_{600}$ of 0.3 followed by 100-fold dilution. One mL of each diluted culture was aliquoted into individual wells of a 96-well culture block and bacteria were grown shaking at room temperature for another 3.5 hours to reach mid-exponential phase. Twenty µL of phage lysate was added to each well at varying concentrations (ranging from 0.001 to 10 phages/bacteria on average) and staggered in time to achieve an adsorption time of 30 minutes (Fig. S5). After allowing phages to adsorb, 200 µL of the phage and bacteria mixture was filtered using a 96-well filter system (Millipore MultiScreen Vacuum Manifold) to remove bacteria and any infecting or adsorbed phages. Five µL of a ten-fold dilution series of each well was then drop spotted onto a fresh lawn of a sensitive host (orange strain 10N.261.55.C8 was used for experiments with 1.143.O and 10N.286.54.F7 was used for experiments with 1.281.O) made in rectangular petri dishes (1-well Nunc Rectangular Dishes, Polystyrene, Sterile by Thermo Scientific (Supplier No. 267060)). Plates were incubated at room temperature for 18-24 hours and then imaged using a flatbed scanner as described above (Fig. S5). Phage adsorption was estimated by comparing the number of plaque forming units (PFUs) in each dilution series to the no host control. For example, in Figure S5A the same order of magnitude of PFUs is evident in both the outgroup and the no-host control, meaning there is no phage adsorption for the outgroup. Yet, there are an order of magnitude more PFUs in both controls compared to the “purple” and “orange” hosts, implying equal adsorption is seen on the “purple” and “orange” hosts.

Bacterial strain selection for transposon mutagenesis and gene deletions

Because the adsorption assays indicated that both orange and purple phages adsorbed to both host groups, we chose one strain, 10N.261.55.C8 (Orange WT, hereafter C8-WT), for mapping of receptors for both host groups. For receptor mapping, we took advantage of the lysis from without phenotype where phages can effect lysis if hosts possess a specific receptor even if no viable phage are produced. Accordingly, at high phage titer, cells of both host groups are lysed by both phages (Fig. S4), allowing for testing of receptors using a "purple" phage on an "orange" host. The same C8-WT strain was used for characterization of resistance determinants of the "orange" host group by gene deletion (see below).

Growth conditions of strains used for transposon mutagenesis and gene deletions

C8-WT was routinely grown at 25ºC in 2216MB or TSB2. The *Escherichia coli* strains were grown in BD Difco Miller Luria-Bertani broth (LB) at 37ºC and supplemented for auxotroph strain *E. coli* Π3813 with thymidine (0.3 mM), and for strains *E. coli* β3914 and MFDpir with diaminopimelic acid (dapA) (0.3 mM). Antibiotics were used at the following concentrations: erythromycin (Erm) 200 µg $mL^{-1}$, kanamycin (Km) 50 µg $mL^{-1}$ and chloramphenicol (Cm) at 5 or 25 µg $mL^{-1}$ for *Vibrio* and *E. coli*, respectively.

Receptor identification using transposon mutagenesis

To map phage receptors, transposon mutagenesis was carried out using suicide delivery of a mariner transposon. C8-WT served as recipient and the dapA deficient strain *E. coli* MFDpir as donor with the suicide conjugative plasmid pSC189-Cm (*61*) (Table S5). The delivery plasmid (pSC189) can be mobilized via RP4-mediated transfer and it carries the hyperactive C9 mariner transposase (*62*). Conjugation was carried out by mating assays as described previously (*63*) with some modifications. First, donor : recipient ratio was adjusted to 1:3. Overnight cultures were diluted 1:100 in fresh media and grown up to an $OD_{600}$ of ~0.4. One mL was separately pelleted at 5,500 x *g* for 2 minutes and washed in pre-warmed mating media broth MMB-1 (TSB supplemented with 1% NaCl plus dapA) to remove antibiotics and/or residual media. This wash step was repeated twice. Washed pellets were subsequently mixed in the same tube with 500 µL of MMB-1, pelleted and resuspended in a mating spot (20 µL) on mating media agar plates and incubated at 25ºC for 18 hours. Mating spots were collected using a Nunc 10 µL sterile plastic inoculation loop and resuspended in 500 µL of ASW. Then, 100 µL of this suspension were spread onto TSB2 plates supplemented with Cm and incubated at 25ºC for 48 hours. Finally, the mutant library (totaling 26,662 mutants) was archived in 500 µL aliquots with ASW supplemented with Cm and 25% glycerol (v/v), quickly frozen in a dry ice bath for 10 minutes, and stored at -80ºC until testing.

Resistant mutants were selected by challenging the mutant library with high titers of phages. Four aliquots of the mutant library were defrosted, centrifuged by pelleting at 5,000 x *g* for 5 minutes, and then washed with 2216MB to remove any residual glycerol. This wash step was repeated twice, and the washed pellets were then resuspended in their original tube with 1 mL of fresh 2216MB supplemented with Cm. C8-WT served as positive control and was treated equivalently, except for the addition of Cm. The washed mutant library and C8-WT control were both diluted 1:10 in 2216MB and then incubated at room temperature with vigorous shaking (250 rpm) for 1 hour until the cultures reached early exponential phase. To select for phage-resistant mutants, lysates was serially diluted 10-fold and mixed with the mutant library and C8-WT cultures. Aliquots of host-phage culture were mixed into 750 µL of 2216MB top agar (with and without Cm as needed) and spread on large 2216MB bottom agar plates (with and without Cm as needed) following the soft agar protocol as described previously (*39*). After incubating at room temperature for 48 hours, ~100 phage-resistant colonies were selected at random and serially re-streaked three times on 2216MB agar plates (with and without Cm as needed). Glycerol stocks of each mutant were archived, and all mutants were then re-tested for phage susceptibility. The re-test was always done with two “orange” and two “purple” phages, one of each always being the original phage used to isolate resistant colonies. In all cases, resistance to one “orange” phage yielded resistance to all “orange” phages and resistance to one “purple” phage yielded resistance to all “purple” phages. Cross resistance to opposite or both phage groups was never seen, further supporting the finding that each group of phages uses a different receptor.

Arbitrary PCR (*64*) was used to map the transposon insertions in resistant strains. Genomic DNA from each phage-resistant mutant was extracted with Lyse-n-Go direct PCR reagent (Thermo Scientific), and 1 µL of the lysate served as template in arbitrary PCR. This method involved two rounds of PCR amplification (*64*): in the first round, genomic DNA was amplified with a fully degenerate primer SS9arb2 (Table S6) containing a 5´ tail of known sequence to be used for specific amplification in the second round of PCR (*65*), paired with primer Mar4 (Table S6) that binds the end of the transposon TnSC189 (*66*). Optimized conditions for the first round PCR consisted of the following reagent concentrations and amplification parameters: primers SS9arb2 and Mar4 were at 0.5 mM and 0.2 mM, respectively; GoTaq G2 HotStart (Promega) was used with $MgCl_2$ at 2 mM; initial heating for 2 minutes at 95°C, followed by 6 cycles of 30 seconds at 95°C, 30 seconds at 30°C, and 1 minute and 30 seconds at 72°C; 30 cycles of 30 seconds at 95°C, 30 seconds at 55°C and 1 minute and 30 seconds at 72°C, with a final extension for 5 minutes at 72ºC. In the second round of PCR amplification, 2.5 µL of the first-round PCR product was used as template, combined with a nested primer within the amplified fragment of TnSC189 (Mar4_int2) and primer (Arb3) with sequence identity to the 5' tail of the SS9arb2 (Table S6). For the second round, PCR reagents were used as described above but using both primer concentrations were 0.2 mM, and the PCR was run under the following conditions: 2 minutes at 95°C, 30 cycles of 30 seconds at 95°C, 30 seconds at 58°C and 1 minute and 30 seconds at 72°C, with a final extension time of 5 minutes at 72ºC. PCR products were verified by electrophoresis, purified by spin-column using QIAquick PCR Purification (Qiagen) and then Sanger-sequenced. Finally, amplicons were trimmed and mapped to the C8-WT genome to identify transposon insertion locations using a custom python script and BLASTn (*67*). Hits are present in Table S2.

<u>Receptor verification using re-sequencing of spontaneously resistant isolates</u>

As an independent method to transposon mutagenesis, to identify phage receptors, we re-sequenced spontaneously resistant mutants from co-cultures of orange host C8-WT and high titer phages. C8-WT was streaked out from glycerol stock onto 2216MB agar plates, inoculated into 5 mL of 2216MB liquid media, gown shaking overnight at room temperature, and plated as a lawn in a soft agar overlay. Five µL drop spots of a phage dilution series were plated on top of the agar, and after 24 hours, resistant colonies that grew in the presence of high phage concentrations were re-streaked three times and archived. For each phage, 10 colonies were archived. Resistant strains were re-streaked and re-tested to verify resistance. The re-test was always done with two "orange" and two "purple" phages, one of each always being the original phage used to isolate resistant colonies just as in the transposon experiments. The results were also consistent with the transposon mutagenesis experiments: In all cases, resistance to one "orange" phage yielded resistance to all "orange" phages and resistance to one "purple" phage yielded resistance to all "purple" phages. Cross resistance to opposite or both phage groups was never seen, further supporting the finding that each group of phages uses a different receptor. Six to seven strains verified in this way were sequenced on an Illumina HiSeq as described in the "Hybrid genome assemblies" section. Reads were trimmed and mapped to the hybrid assembly reference genome using CLC work bench 9. Single nucleotide polymorphisms (SNPs) and indels were identified using a custom pipeline made for CLC work bench 9 and are presented in Table S3. These SNPs were cross-referenced to receptor identification using transposon mutagenesis (see above).

Identification and annotation of putative PDEs in the flexible genome

In order to determine the differences in the flexible genome amongst the 19 "orange" and "purple" strains, we created a multiple alignment using Mugsy (*68*), and performed a hierarchical clustering of the alignment blocks, greater than 500 bp, by length in R using hclust (*69*) (Fig.1A, right). The two groups clustered by their phage predation profile. This clustering was completely driven by the presence of 5 alignment blocks, three of which were exclusive to the "orange" strains, and two of which were exclusive to the "purple" strains. Upon further investigation of the alignment blocks by hand, we discovered that the alignment blocks corresponded to putative PDEs (Fig. 1B).

Gene annotations of the PDEs were performed manually using the consensus obtained from HHPred (*70*), InterProScan5 (*58*), Phyre2 (*45*), and BLASTp (*71*) databases tools (Data S1). The search was performed using default options except that HHPred search was performed against COG-KOG 1.0 and Pfam-A_v32.0 databases and that BLASTp was performed using the protein-protein BLAST option. Up to ten significant pfam and COG ($p < 0.05$) from HHPred search where used to compare each gene with pfam-COG accession numbers of phage defense systems from supplementary Table 1 from Doron et al. (*16*).

To further characterize the distribution of the PDEs identified among the 19 strains in a larger collection of *Vibrio* genomes (*32*), we used BLASTn (*67*) and custom python scripts to identify the distribution of the mobile elements (Fig. S6). Because when comparing long mobile elements BLASTn will often return multiple overlapping ranges of identity, our BLAST parsing script merges overlapping sequence ranges to avoid over-counting regions within a genome. A PDE was considered present in a genome if at least 80% of the element was present at over 95% identity.

Methylation profiling

In order to discover if the restriction modification systems identified on the PDEs in the "orange" and "purple" strains are active, we determined the methylation sites in both phage-host pairs as outlined in Murray et al. (*72*). Briefly, we submitted the host genomes to REBASE, a well-curated database of restriction modification systems which allows motif prediction based on comparisons to known enzyme-motif pairs (*73*). Then, we combined the motif data with the methylome data generated using the Base Modification Detection and Motif Analysis pipelines available on the single molecule real-time (SMRT) sequencing portal from Pacific Biosciences. Summary data is presented in Table S4.

Phage defense element knockouts using two-step allelic exchange

To test whether putative PDEs were responsible for phage resistance in C8-WT, we knocked out large portions of each PDE containing genes annotated as being related to phage defense (Fig. S6), in all possible combinations. We found it was possible to knock out nearly all of PDE1 (93.5%), but had to leave in part of the element's Yfbr gene as it replaces the host Yfbr gene upon insertion. For PDE2, we found that knocking out the entire element was not possible in a single step, likely because of the toxicity effects of deleting the entire TA system at once. We therefore proceeded to make a partial deletion (58.8%) from the toxin gene to the 5' end of the element, leaving the antitoxin intact, along with other genes predicted to play roles in insertion/mobilization of the element (integrases, transposases, and recombinases) as indicated by structure and function annotations using HHpred (*74*), InterProScan5 (*58*), and Phyre2 (*45*) (Fig. S6, Data S1). Noting PDE3 also has a putative TA system, we followed the same approach as for PDE2 and made a partial deletion (68.8%) from the toxin gene to the 5' end of the element. The details of the approach are summarized in Figure S6.

Site-directed mutagenesis was used for all deletions. Cloning was carried out using the New England Biolabs Gibson Assembly Master Mix according to the manufacturer's protocol. Fragments upstream and downstream of the portion of the element to be deleted were separately PCR-amplified using primers specified in Table S6. A third PCR reaction was carried out to amplify the backbone of the plasmid pSW7848T (*75*) (Table S5) with primer pairs pSW_F&R

(Table S6). These amplicons were cut with Dpn1 (2 hours at 37ºC) to inactivate the plasmid template before setting up the Gibson assembly reaction. In all cases, PCR products were verified by gel electrophoresis, purified by spin-column as described above, and DNA concentration was determined using Nanodrop 2000 (Thermo Scientific). Subsequently, 0.03 pmol of each vector was assembled with 0.07 pmol of its specific downstream and upstream DNA fragments at 50ºC for 60 minutes. After completion of this reaction, DNA was desalted by dialysis on a 0.0025 µM filter (Millipore) before electroporation into *E. coli* Π3813, which was used as a plasmid host for cloning (*63*). Finally, the plasmid DNA was purified, verified by Sanger sequencing, and electroporated into *E. coli* β3914 to be uses as a plasmid host for conjugation (*63*) (Table S5).

Conjugation was carried out in a mating spot as described above for the transposon mutagenesis but with some modifications: donor : recipient ratio was changed to 3:1, the mating media was altered to MMB-2 (TSB supplemented with 2% NaCl plus dapA), and the mating spot was incubated at 30ºC. Counter-selection of ΔdapA donor was performed by plating on TSB2 agar plates without dapA but supplemented with Cm and glucose 1% (w/v). Antibiotic-resistant colonies are due to the integration of the entire plasmid ($Cm^R$) in the chromosome by a single crossover. Colonies were picked, re-grown in liquid media (TSB2) supplemented with Cm and glucose 1% (w/v) to late logarithmic phase and spread on BD Bacto TSB without Dextrose plates supplemented with 2% NaCl (w/v) and 0.2% arabinose. To verify deletions in the single PDE mutants ΔPDE1, ΔPDE2 and ΔPDE3 (Table S5) PCR products generated using primers flanking externally the different regions targeted (ΔPDE1/F&R; ΔPDE2/F&R and ΔPDE3/F&R) (Table S6) were sequenced by Sanger. This procedure was also used to construct double (ΔΔPDE12; ΔΔPDE13; ΔΔPDE23) and triple mutants (ΔΔΔPDE123) but using a single or double mutant as final recipient during the conjugation step (Table S5).

<u>Phage susceptibility assay for mutants</u>

In order to test the susceptibility of the "orange" PDE deletion mutants to "purple" phages, we challenged each mutant with representative "purple" phage 1.281.O in agar overlays (Fig. 2) and in liquid culture (Fig. S7). For the mutant testing in agar overlays, we used the same protocol outlined in the "Host range assays at varying phage concentrations" section above, with one additional step: after the plaques were imaged, we re-streaked the phages from the highest concentration drop spot onto fresh bacterial (host or mutant) lawns to test for phage propagation (Fig. 2B). For the liquid assay, we streaked out each mutant onto 2216MB agar plates, allowed 48 hours for large colonies to form, and inoculated each mutant into 3 mL of 2216MB in triplicate. After growing the cultures shaking at 25°C for 4 hours, we normalized 1 mL of the culture to an $OD_{600}$ of 0.3, diluted it 100x into a final volume of 5 mL, and aliquoted 200 µL into 12 wells each of a 96-well clear bottom Micro-titer plate (Falcon). A Tecan Microplate Reader with Spark software was used to maintain the cultures shaking at 25°C, monitoring $OD_{600}$ every 15 minutes. Once $OD_{600}$ reached 0.3, "purple" phage 1.281.O was added in triplicate at different concentrations to reach the desired multiplicities of infection (Fig. S7B), after which the cultures were returned to the plate reader for the remainder of a 24 hour run. This experiment was run with two mutants, C8-WT, and "purple" host 10N.286.54.F7 each time until all mutants had been tested.

Identification of putative PDEs from comparison of closely related genomes of different bacteria

We extended the search for novel putative PDEs by searching identify nearly clonal genomes of *Vibrio*, *Salmonella*, and *Listeria*. For the 23 *Vibrio* strains, we selected only those with identical ribosomal proteins. For *Salmonella* and *Listeria*, we selected genomes within the same ribotype using the ribosomal MLST database (https://pubmlst.org/rmlst/), filtered to only include NCBI assemblies, and downloaded genomes from ribotypes with more than 20 members. We used ribotype 8354 for *Salmonella* and MLST strain type 5 for *Listeria* to assay putative PDE distribution in an exemplary manner. In all three cases, the final set of genomes was run though a custom kmer-based comparative genomic pipeline to identify flexible regions. All pairs of genomes were compared. First, each genome was split into 31-mers using Jellyfish (*76*), then shared kmers between the genomes being compared were removed and only unique kmers were mapped back to the reference genomes they originated from using Bowtie2 (*77*). Any unique region greater than 1,000 bp was kept and a gap of 3,000 bp was allotted to account for genes that may have been shared between the two genomes splitting a complete region. Regions were checked for duplication and the largest region of any overlapping regions was saved. We then used Mash (*78*) to compare all the unique regions to each other and clustered any region greater than 5kbp with minimum Jaccard similarity of 0.95. We visualized the clustering using Gephi (*79*) and then chose one representative from each cluster by hand to make a final list of unique regions. We then used BLASTn (*67*) and custom python scripts to determine which genomes harbored which elements with >95% identity and >80% length considered a match. We removed any element that appeared in all genomes. Finally, we used HMMER (*41*) to search each element for known defense genes using Supplementary Table 1 in Doron et al. (*46*). Any unique region with one or more hits was considered to be a putative PDE and depicted in Figure S8, Figure S10, and Figure S11 for *Vibrio*, *Listeria*, and *Salmonella*, respectively.

#### Proportion of known defense genes in flexible genome across diverse *Vibrio*

To determine the proportion of known defense genes in other *Vibrio* species, we used the species and population designations from Arevalo et al., (*32*). We based our identification of flexible genes on the method described in Arevalo et al. ORFs were identified with Prodigal 2.6 (*57*) and orthologous genes were clustered using MMseqs2 (*49*). Flexible genes for a given population were defined as orthologs which were present in at least one member but not present in all members of the population. Flexible genes from each genome were then used as a database which we searched for known defense genes using Supplementary Table 1 in Doron et al. (*16*) using HMMER (*41*). Total length of all flexible genes summed for each species and the proportion of genes (by length) with a hit to a known defense gene is shown in Figure S9.

#### Distribution of putative receptors across diverse vibrios

To test how diverse receptor genes identified by mutant analysis in *V. lentus* are across other vibrios, we used the species and population designations from Arevalo et al., (*32*). In cases where a species was composed of multiple populations, we chose to only analyze the population with the most members. We based our identification of core genes on the method described in Arevalo et al. (2019). ORFs were identified with Prodigal 2.6 (*57*) and orthologous genes were clustered using MMseqs2 (*49*). Core genes for a given population were defined as orthologs which were present in a single copy in all members of that population. We aligned all genes within orthologous clusters using MUSCLE (*80*) and calculated the amino acid diversity between each pair of genes as the number of positions with non-identical amino acids divided by the alignment length. Average pairwise amino acid diversity was obtained by taking the mean diversity across all pairs of genes within an orthologous cluster.

The amino acid sequences of each receptor gene were aligned using mafft-linsi (*42*) and average pairwise amino acid diversity was calculated as described above.

Phylogenetic relationships among receptors identified using genetic approaches were determined across a collection of diverse *Vibrio* isolates (40). Receptor orthologues were identified using BLASTn (*71*) with evalue of 1E-8, aligned using MAFFT (*42*), and final trees were constructed using RAxML (*43*) with model GTRCAT.

To estimate whether diverse isolates are decorated with similar lipopolysaccharides (LPS), we extracted the genome region bounded by *gmp* and *gmhD* (*81*), which encodes genes for LPS synthesis. We searched the same *Vibrio* genomes from Arevalo et al. (*32*), that were used for the determination of known defense genes in Figure S9. When the same set of genes was present, we inferred that the LPS structure was highly similar and may thus bind to similar phage receptors. We only analyzed genomes that had LPS regions that were present on a single contig to avoid ambiguity in presence/absence patterns. Genes were clustered using MMseqs2 (*49*) at 90% identity and required to be present in at least 2 genomes. Presence/absence profiles are shown in Figure S12C using hierarchical clustering to order the genomes (rows) and LPS genes (columns). Genes with identical presence absence patterns in Figure S12C have only one tree branch going into the left-most gene (top tree).

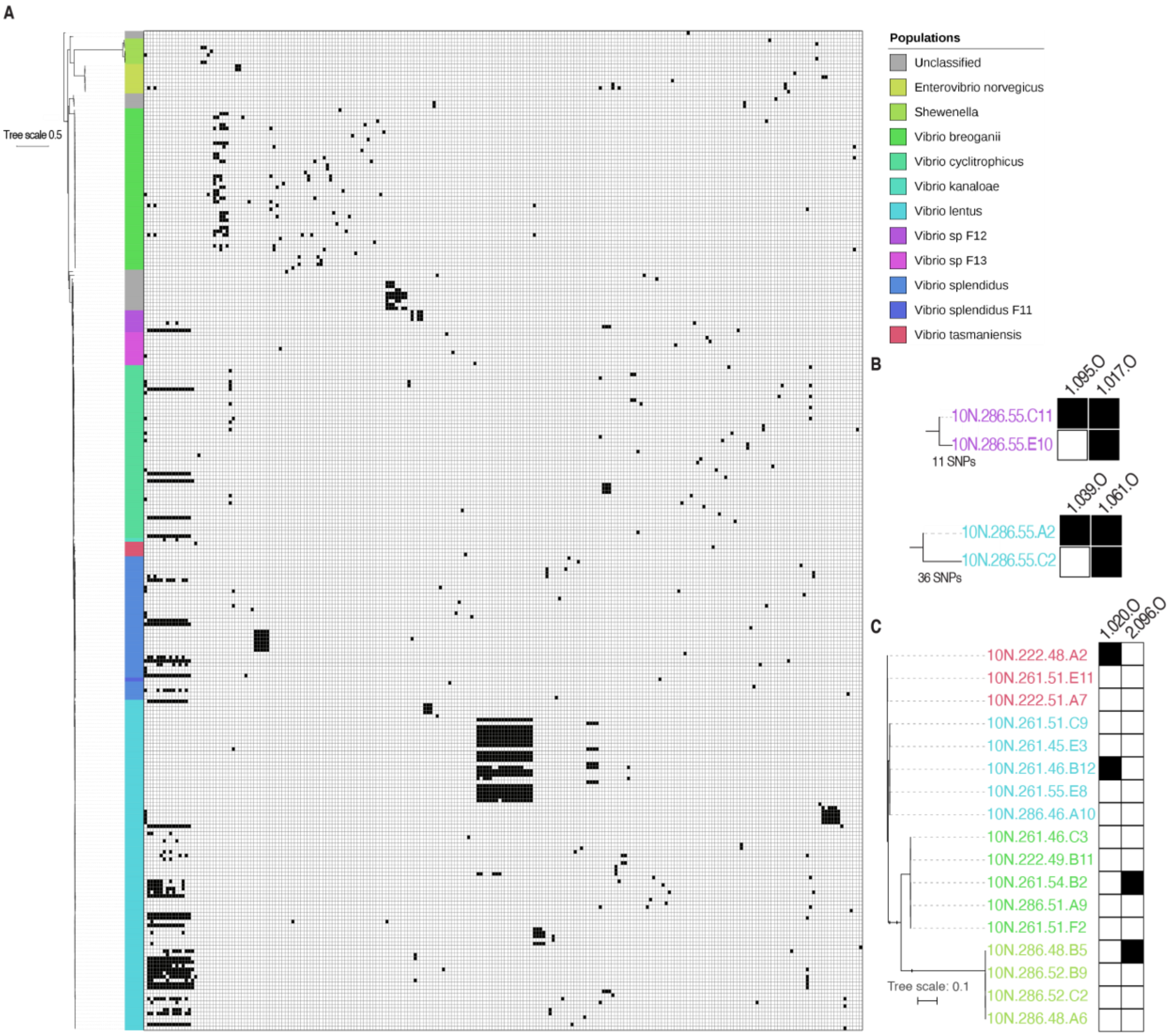


**Fig. S1.**

Phage host-range established using an exhaustive cross-test matrix. **(A)** Full matrix with rows depicting bacterial hosts organized by the phylogeny of their ribosomal protein and *hsp60* gene sequences (proxy for core genome), and columns depicting phages ordered by protein similarity identity [modified from Figure 2 in (*23*)]. **(B)** Closest bacterial relatives differing in phage sensitivity profiles can be distinguished by only few SNPs across their entire core genomes. Trees represent full genome alignments, phage identification codes written above columns, black boxes indicate positive infection determined by plaque assay. **(C)** Broad host-range phages, defined as host ranges spanning different species, remain strain-specific within different species. Phylogenetic tree constructed using same alignment of core genes as in A, and infection representation analogous to that in B.

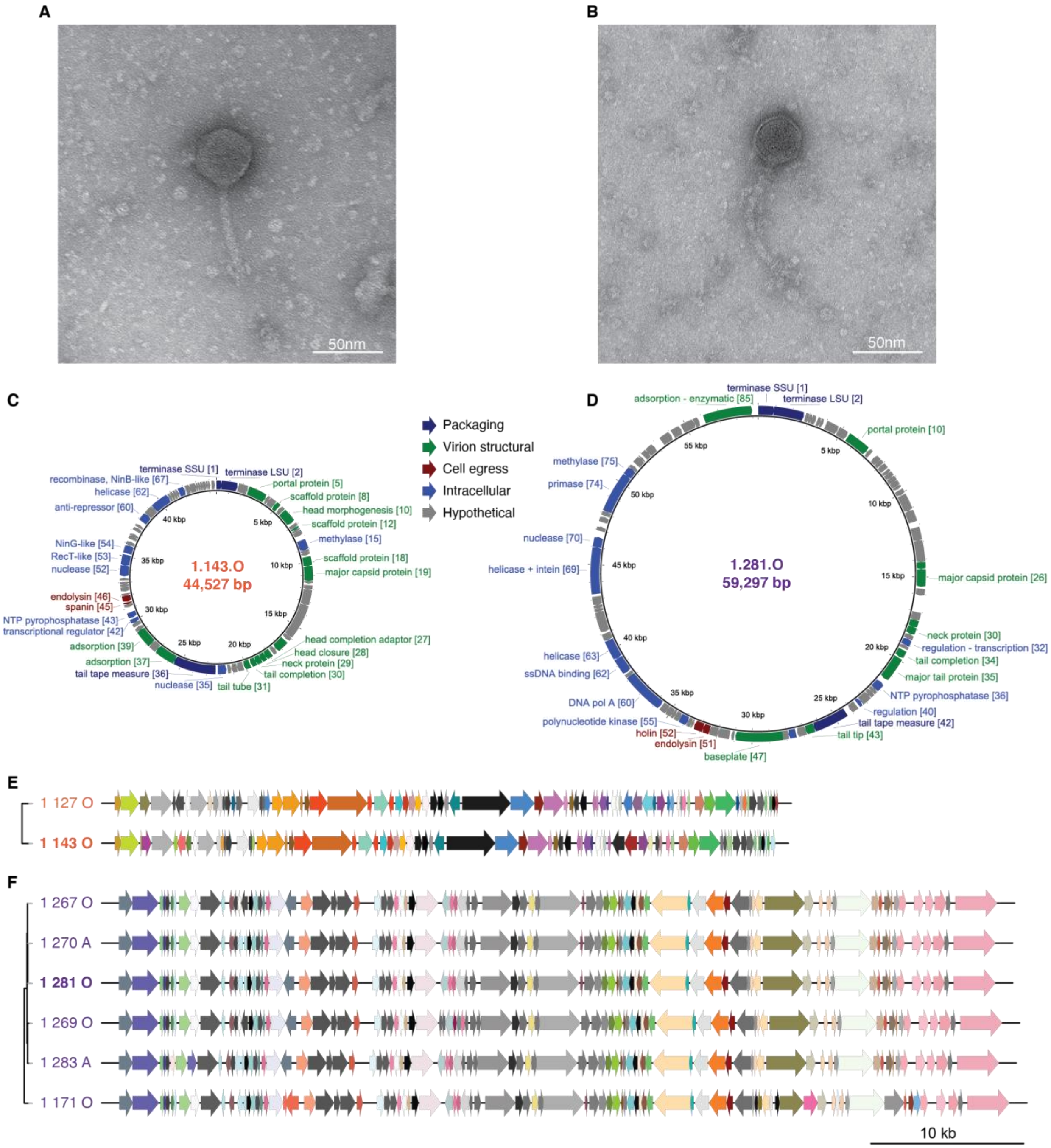


**Fig. S2.**

"Orange" and "purple" phages represent divergent groups of siphoviruses. **(A, B)** Electron microscopy of phages representative of "orange" and "purple" groups suggests that both are siphoviridae, with long non-contractile tails. **(C, D)** Genome characterization of phages representative of "orange" and "purple" groups, respectively, shows that they differ in size by nearly 15 kbp; numbers adjacent to annotations reflect GenBank locus tag. **(E, F)** Clustering and alignment of phage genomes show that they represent two distinct genus-level groupings. While within each group gene synteny and content are conserved, no gene clusters are shared between groups.

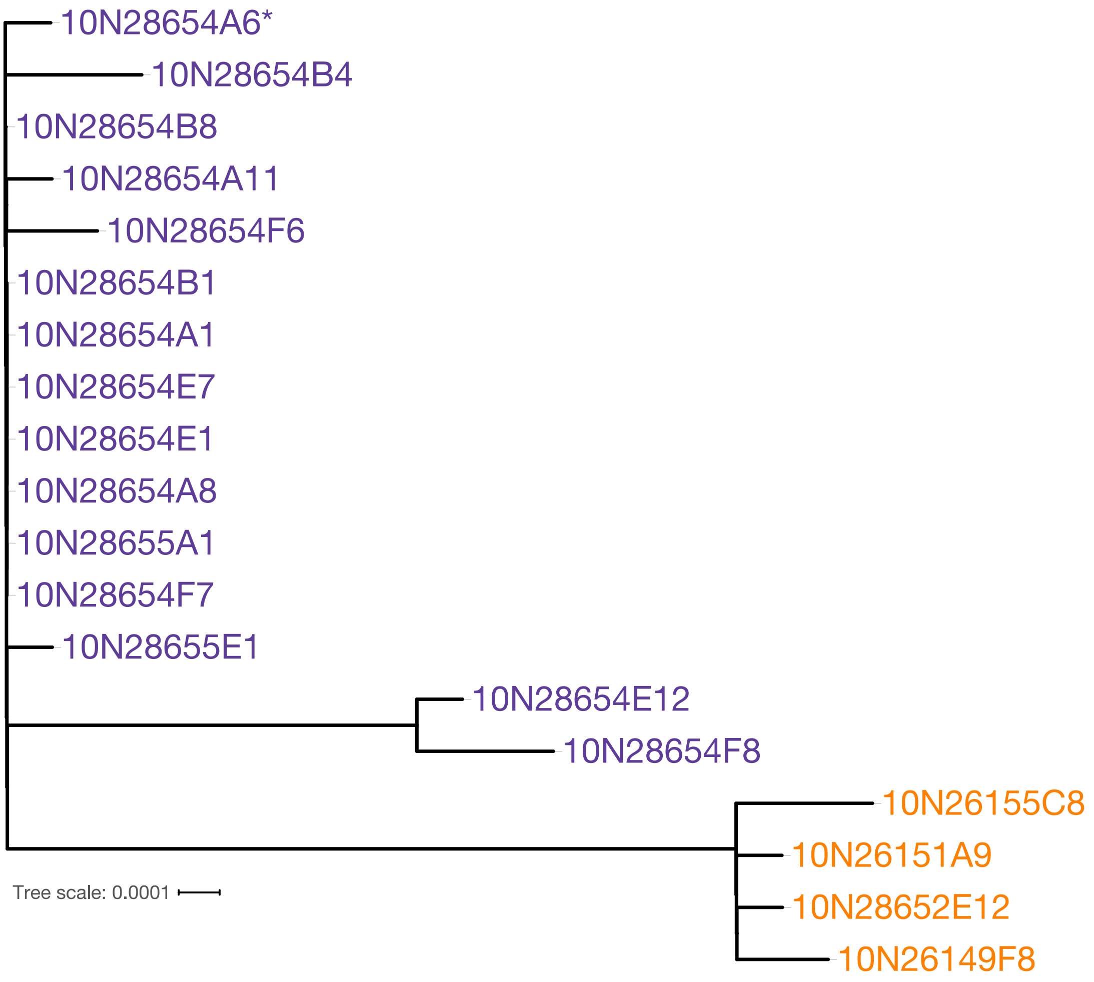


**Fig. S3.**

Unrooted maximum likelihood tree for core genomes of all the nineteen "purple" and "orange" host clones. The strain chosen by the Parsnp program as a reference is indicated by *. 44 SNPs were identified in the total alignment and 14 SNPs differentiate the "orange" and "purple" subsets (see Table S1 for full list).

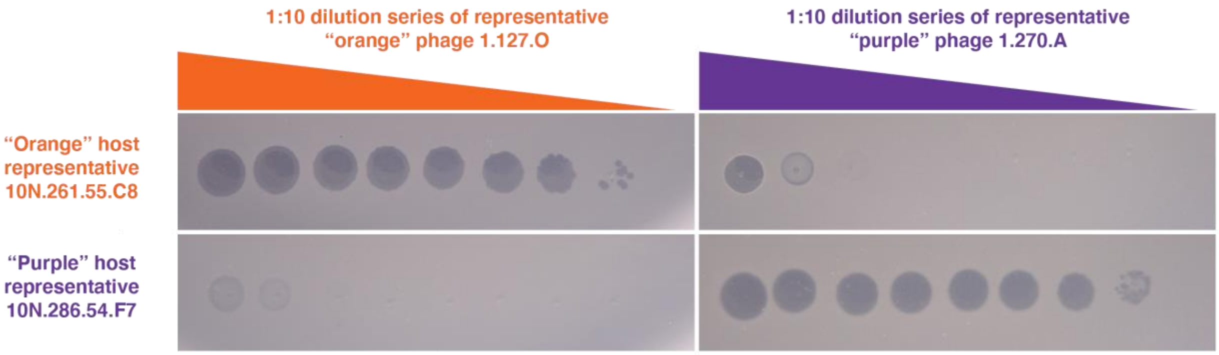


**Fig. S4.**

Efficiency of plating assay demonstrating effect of differing phage concentrations on host killing. At high concentrations, phage can effect lysis even of non-hosts but without production of viable progeny (“lysis from without”) indicating that phage can attach and enter the cell, but that replication is prevented internally.

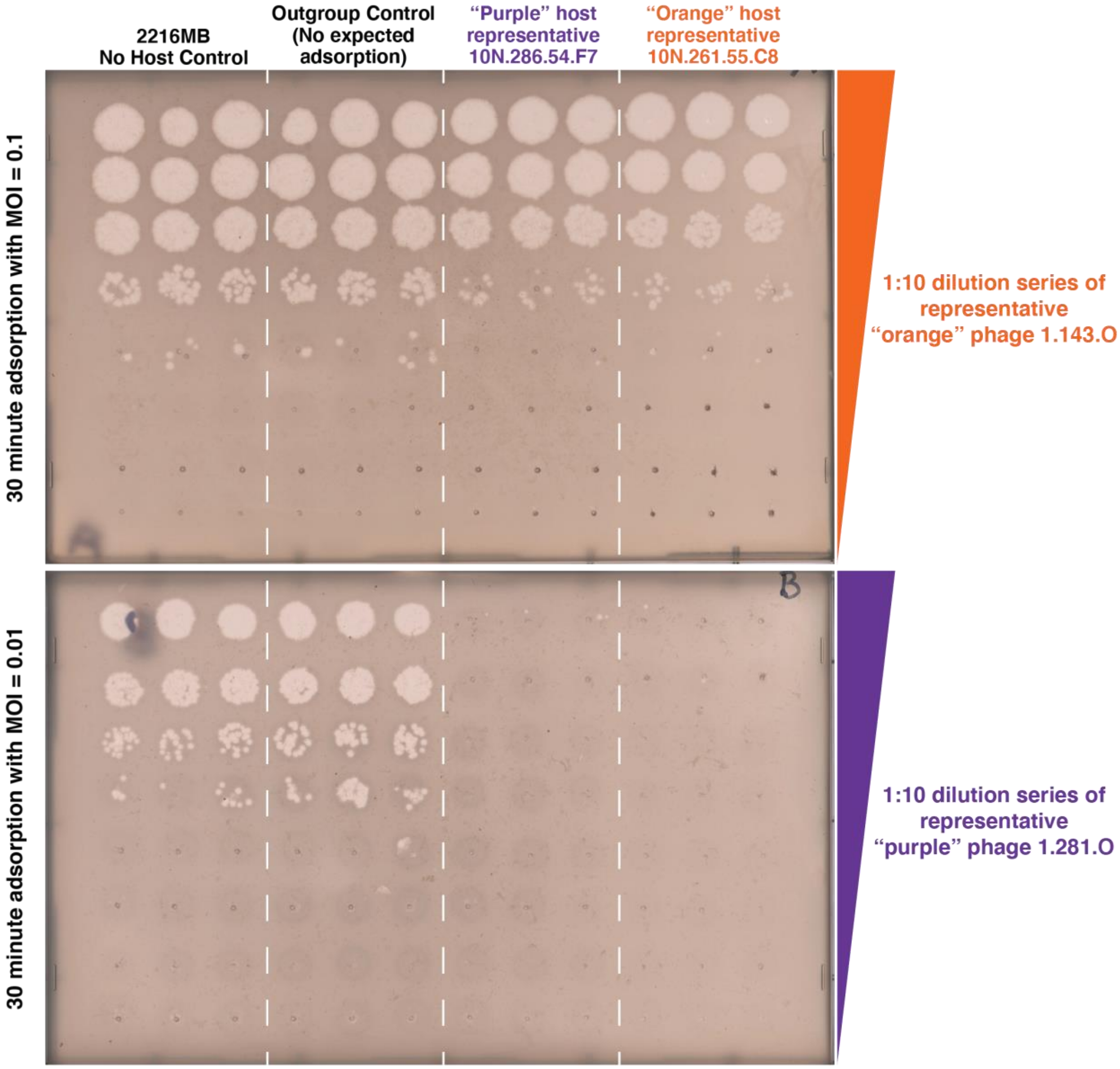


**Fig. S5.**

Phage adsorption assay showing that phage can adsorb to both "orange" and "purple" strains irrespective of whether those bacterial strains can serve as hosts for viable phage production. After allowing a fixed concentration of phages to adsorb to different bacterial strains, free phages that remained unattached were plated with sensitive hosts to quantify adsorption as the difference to no-host controls (see methods). Both "orange" and "purple" phages were found to adsorb to "orange" and "purple" hosts, but not to an outgroup control. In the top panel, "orange" phage 1.143.O shows the same adsorption phenotype to both "orange" host 10N.261.55.C8 and "purple" host 10N.286.54.F7: the number of free phages decreased by ten-fold. In the bottom panel, "purple" phage 1.281.O shows the same adsorption phenotype to both "orange" host 10N.261.55.C8 and "purple" host 10N.286.54.F7, attaching with full efficiency. In both cases, no attachment is observed for a *Vibrio* outgroup host (10N.261.49.C11) as indicated by the same level of phages as in no host controls.

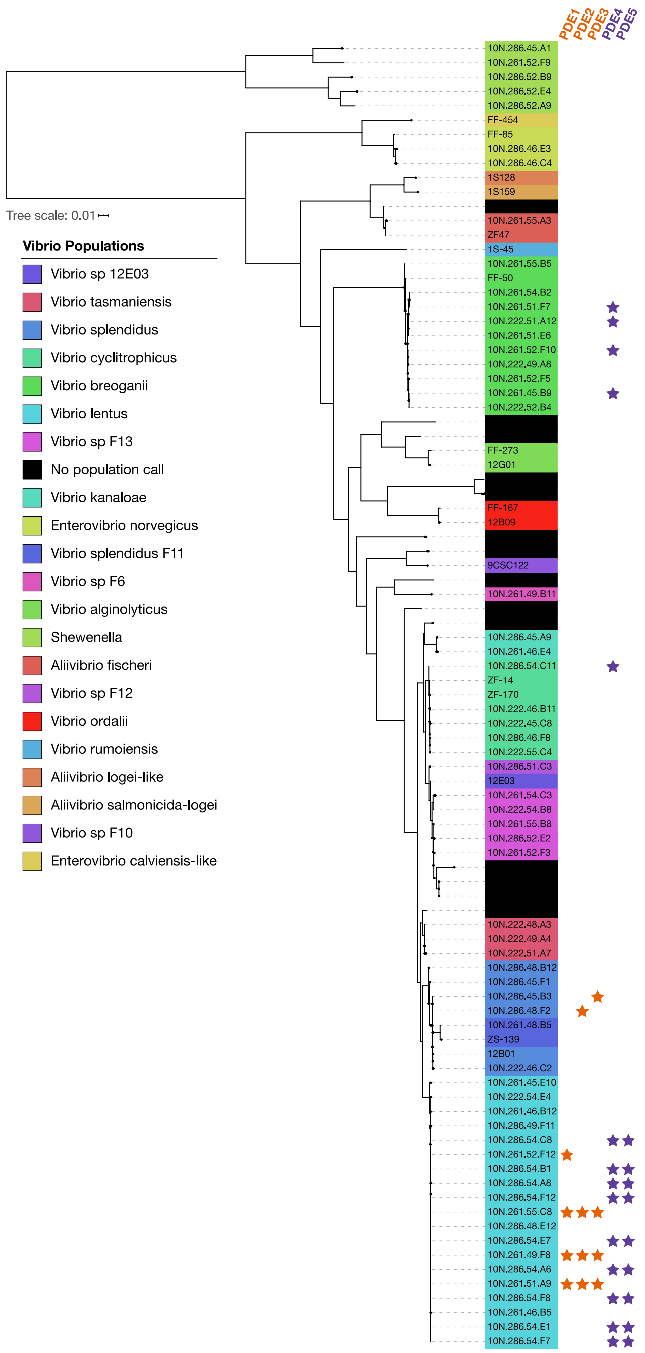


**Fig. S6.**

Presence of the same phage defense elements (>95% nucleotide identity over >90% of the total element length) in divergent genomic backgrounds suggests their movement via horizontal gene transfer. Pruned tree from Figure S1 depicting the phylogeny of ribosomal protein and *hsp60* gene sequences (proxy for core genome) of each *Vibrio* host.

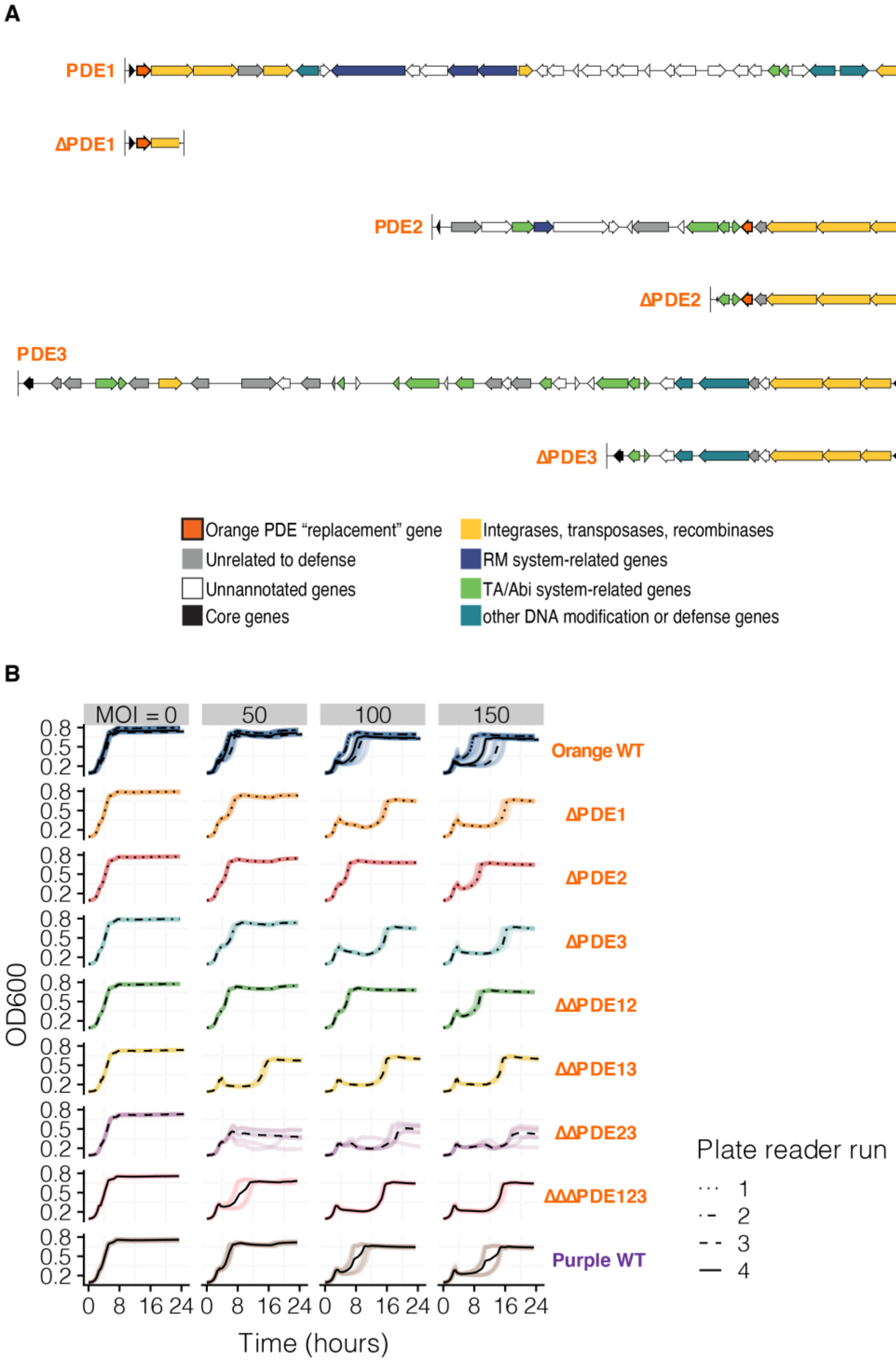


**Fig. S7.**
PDE deletions and phage susceptibility testing. (A) Genetic knockout diagrams for each phage defense element in the “orange” strains, and (B) growth curves of each combination of knockouts grown to mid-exponential phase and then challenged with “purple” phage 1.281.O at varying concentrations.

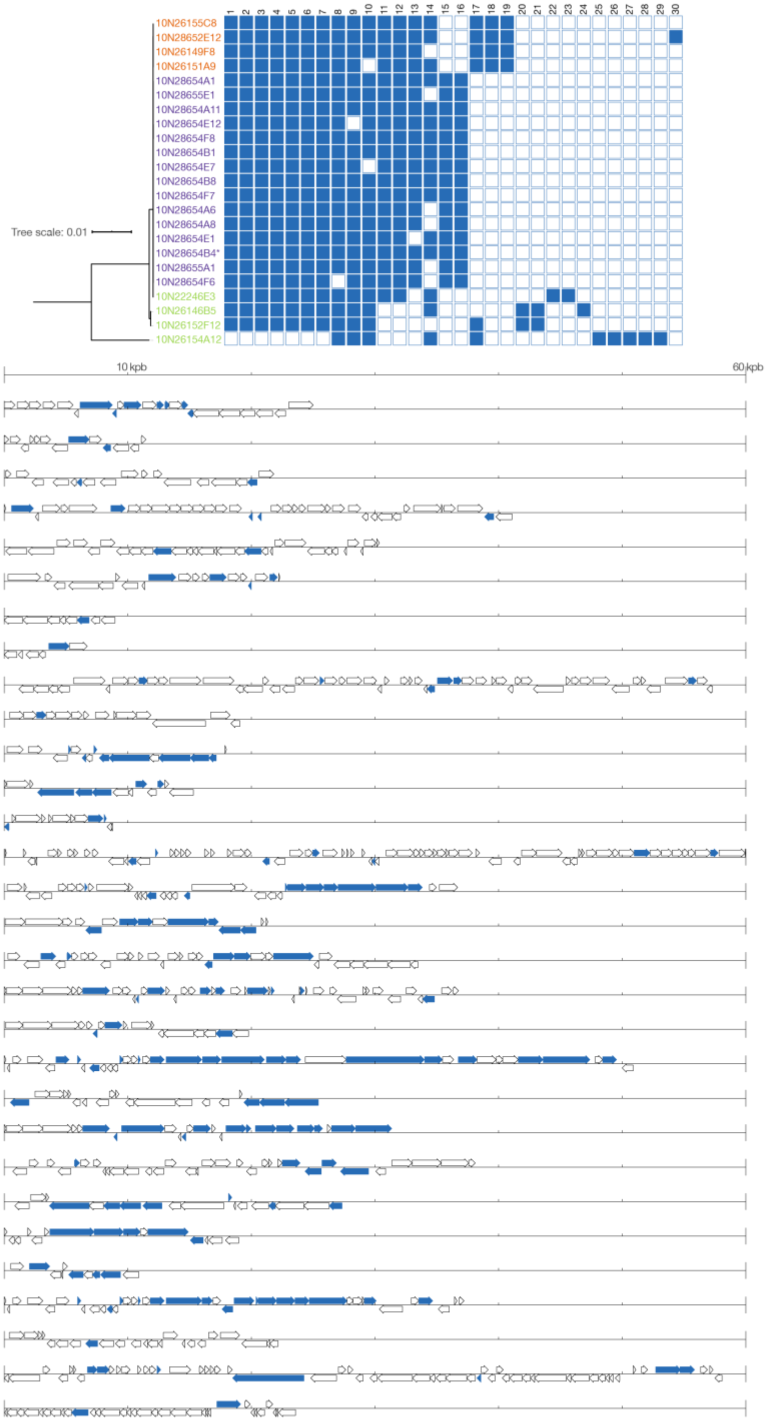


**Fig. S8.**
Distribution of all putative PDEs in *Vibrio lentus* clones with accompanying gene diagrams. Bacterial hosts are arranged by core genome tree. In gene diagrams, hits to known defense genes are shown in blue.

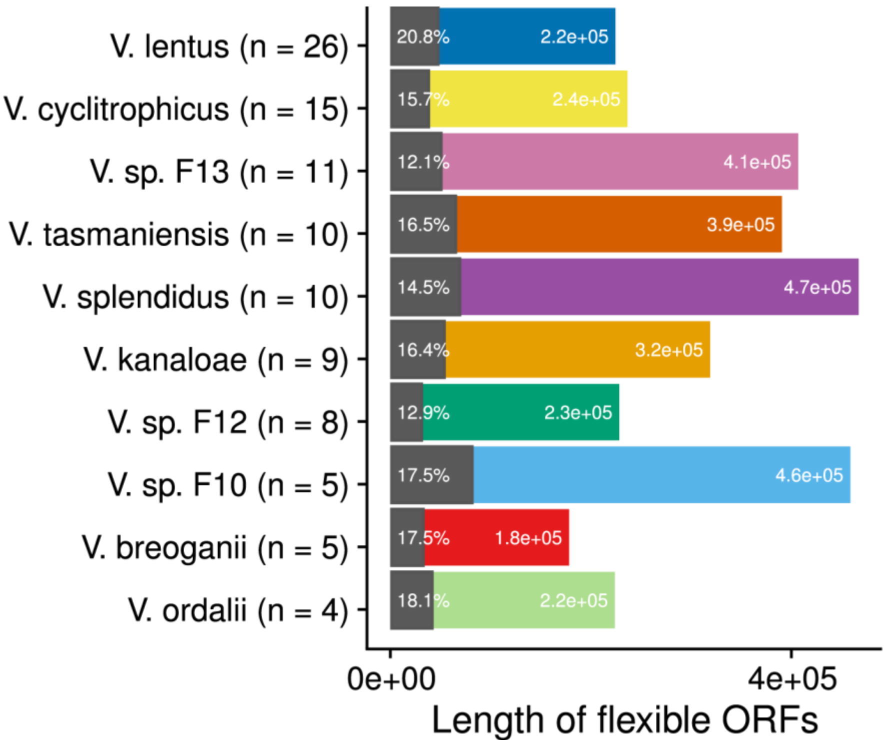


**Fig. S9.**
Proportion of known phage defense genes by length in the flexible genomes of diverse *Vibrio* species. Between 12-21% of the flexible gene content of ten different species, represented as populations defined as gene flow clusters (*32*), can be attributed to known phage defense genes.

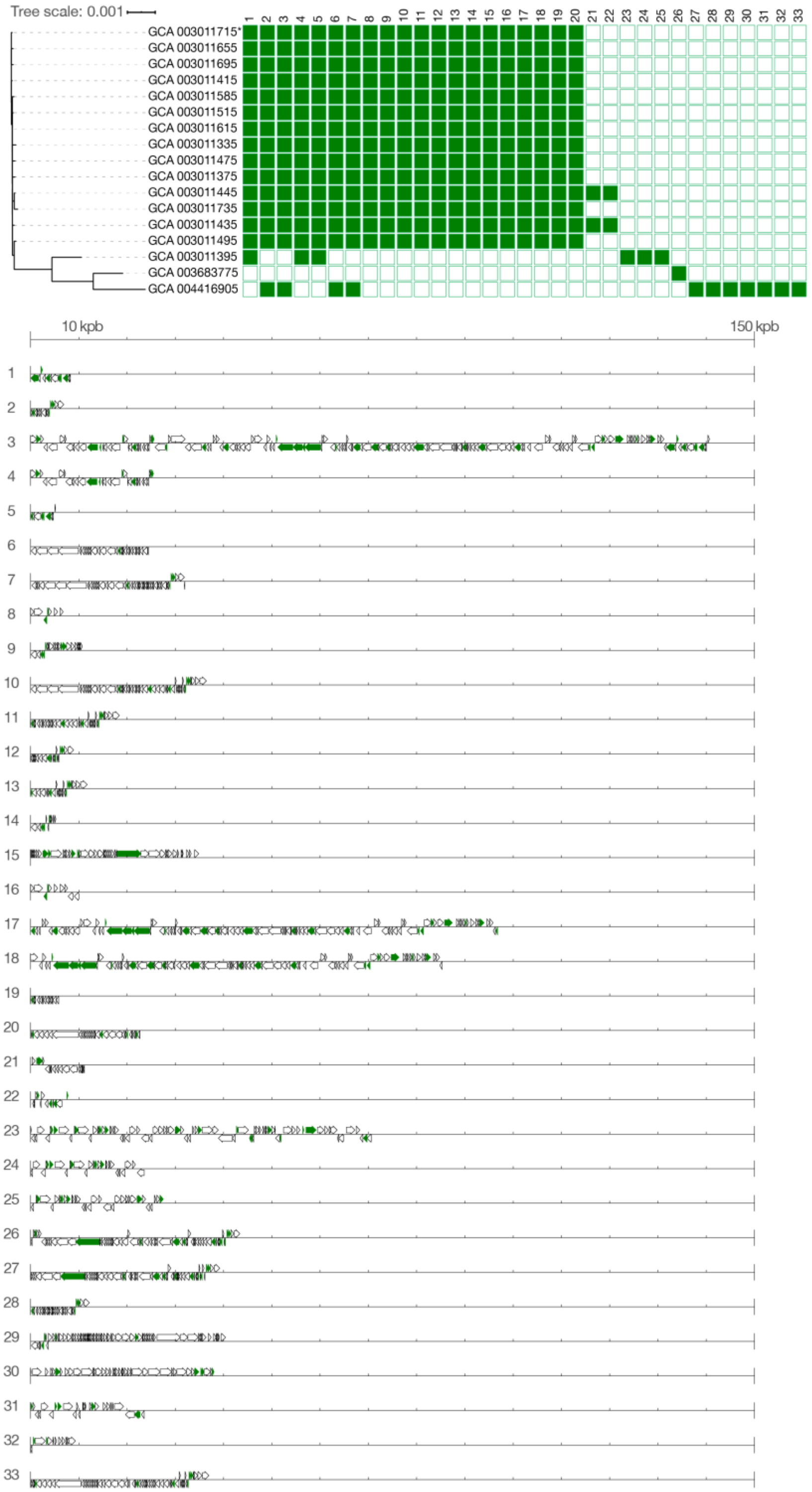


**Fig. S10.**
Distribution of all putative PDEs in *Listeria* genomes with accompanying gene diagrams. Bacterial hosts are arranged by core genome tree. In gene diagrams, hits to known defense genes are shown in green.

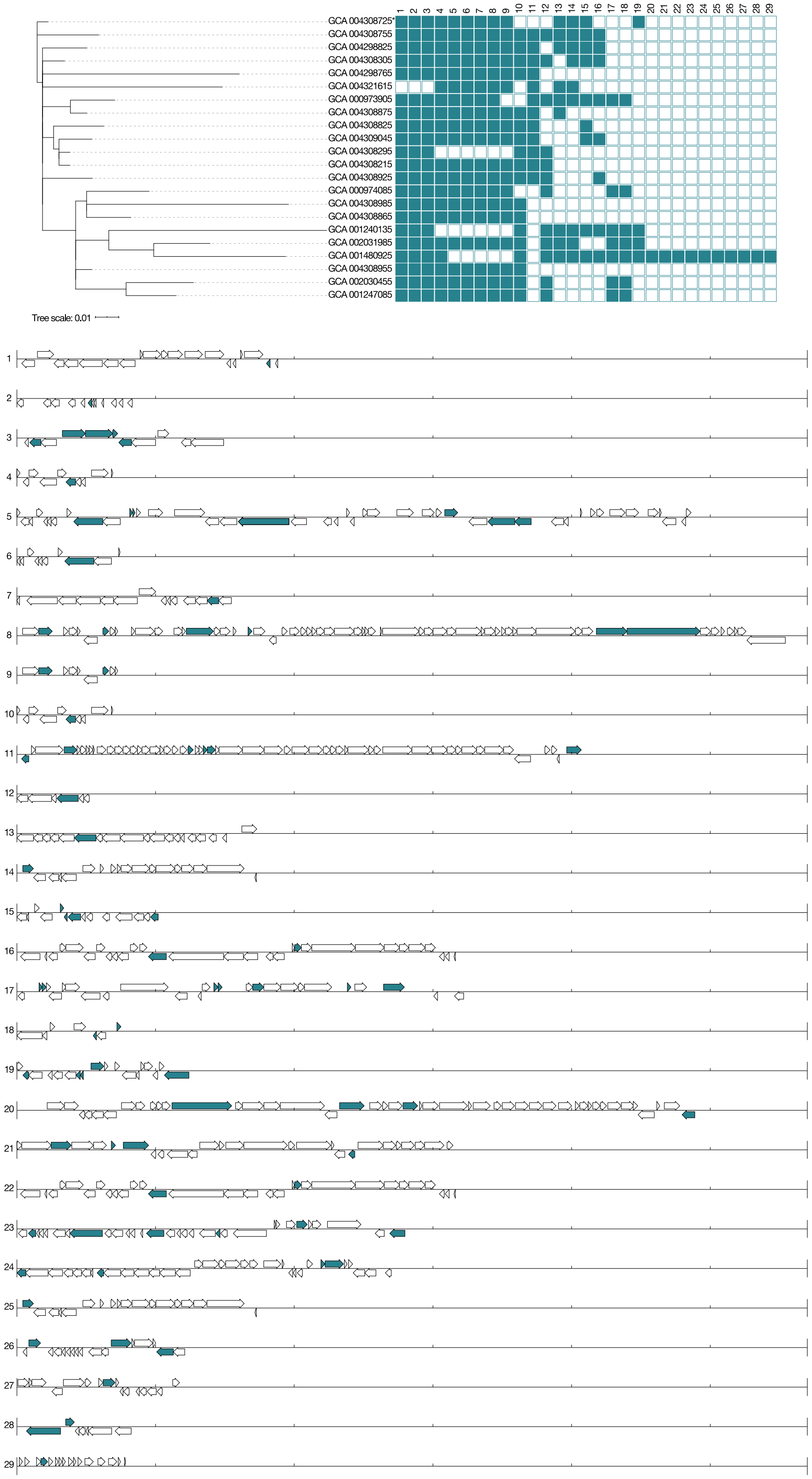


**Fig. S11.**
Distribution of all putative PDEs in *Salmonella* strains with accompanying gene diagrams. Bacterial hosts are arranged by core genome tree. In gene diagrams, hits to known defense genes are shown in teal.

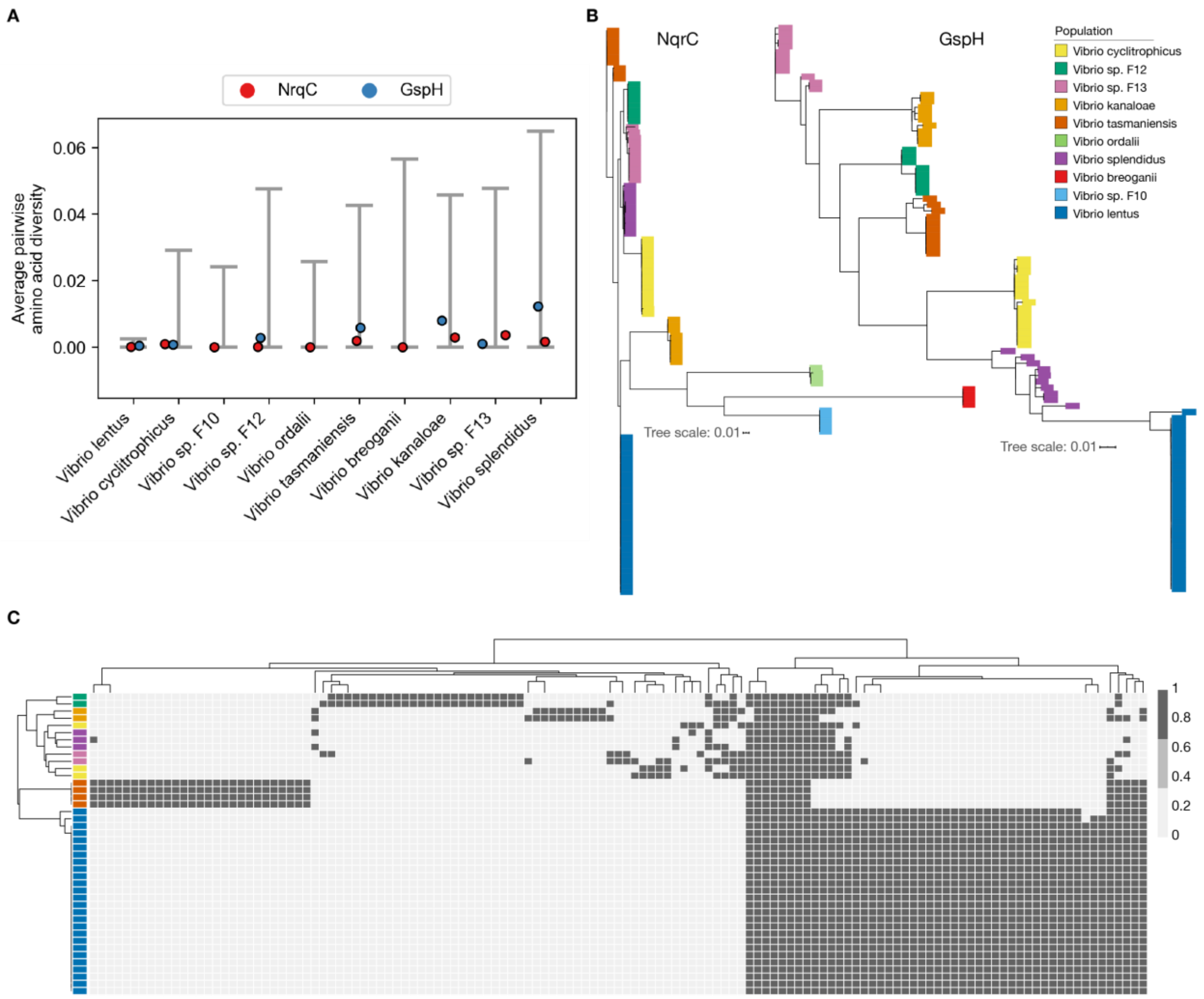


**Fig. S12.**

Phage receptor diversity across diverse *Vibrio* species suggests positive selection at the population level. **(A)** Genes identified as receptors in *V. lentus* show much reduced diversity compared to core genes in 10 populations of vibrios. Populations are defined as co-existing members of species and represent gene flow units (*32*). Each point represents the average amino acid pairwise diversity for a specific receptor gene indicated by color. The grey bars represent the 2.5th and 97.5th percentile range of average amino acid pairwise diversity for all core genes within a given population. **(B)** Nucleotide trees of select receptor genes show evidence of recent population-level sweeps since receptor alleles within each population are nearly identical. The outermost protein of the sodium transporter (NqrC) is most likely to be in physical contact with a phage tail, and the pseudopilin (GspH) had the most transposon insertions in strains resistant to the “orange” phage is the screen (Table S2). **(C)** Genes encoding the LPS show the same presence/absence patterns within populations indicating that they encode the same LPS variant, indicating receptor evolution is congruent with population evolution.

**Table S1.**

SNPs in the core genome of 19 “purple” and “orange” clones (matching Fig. S3).

| SNP # | Nucleotide change P-->O | ORF in 10N28654F7 | Annotation | AA change P-->O | Notes |
|---|---|---|---|---|---|
| 1 | G-->T | ORF_0_1115 | Cytochrome O ubiquinol oxidase subunit III | Val-->Phe | |
| 2 | C-->T | ORF_0_1604 | Unannotated | NA | Asp-->Asp; Near end of a PDE |
| 3 | C-->T | ORF_1_1456 | Pesticidial crystal protein cry6Aa | Gln-->Stop | |
| 4 | G-->A | ORF_1_917 | Aerobic cobaltochelatase CobS subunit | Ala-->Val | |
| 5 | G-->A | ORF_1_912 | TRAP-type C4-dicarboxylate transport system, periplasmic component | NA | Gly-->Gly |
| 6 | A-->G | ORF_1_365-366 | Unannotated | Stop-->Gln | |
| 7 | C-->T | ORF_0_3133 | Phosphoenolpyruvate carboxylase | NA | Gly-->Gly |
| 8 | C-->T | ORF_0_1978 | IcmF-related protein | Ala-->Val | |
| 9 | A-->G | ORF_1_87 | Unannotated | Thr-->Ala | |
| 10 | A-->G | intergenic | NA | NA | upstream of ORF_1_86: putative orphan protein; putative membrane protein |
| 11 | C-->G | ORF_1_687 | RND multidrug efflux transporter; Acriflavin resistance protein | Gln-->Glu | |
| 12 | G-->A | ORF_0_1847 | Formate dehydrogenase -O, gamma subunit | Met-->Ile | |
| 13 | C-->A | intergenic | NA | NA | upstream of ORF_0_138:tRNA uridine 5-carboxymethylaminomethyl modification enzyme GidA |
| 14 | G-->A | ORF_1_1585 | Probable MFS transporter | NA | Ala-->Ala |

**Table S2.**
Receptor identification by transposon mutagenesis.

| Host | Phage | Gene ID | Annotation | Tn Hits |
|---|---|---|---|---|
| 10N26155C8 | 1.143.O | 10N26155C8_2_159 | General secretion pathway protein H | 81 |
| 10N26155C8 | 1.143.O | 10N26155C8_2_164 | General secretion pathway protein C | 9 |
| 10N26155C8 | 1.143.O | 10N26155C8_2_162 | General secretion pathway protein E | 1 |
| 10N26155C8 | 1.143.O | 10N26155C8_2_156 | General secretion pathway protein K | 1 |
| 10N26155C8 | 1.281.O | 10N26155C8_2_94 | dTDP-4-dehydrorhamnose reductase (EC 1.1.1.133) | 58 |
| 10N26155C8 | 1.281.O | 10N26155C8_2_93 | dTDP-4-dehydrorhamnose 3,5-epimerase (EC 5.1.3.13) | 8 |
| 10N26155C8 | 1.281.O | 10N26155C8_2_96 | dTDP-glucose 4,6-dehydratase (EC 4.2.1.46) | 6 |
| 10N26155C8 | 1.281.O | 10N26155C8_0_669 | Phosphoglucomutase (EC 5.4.2.2) | 3 |
| 10N26155C8 | 1.281.O | 10N26155C8_0_485 | ABC-type multidrug transport system, ATPase and permease component | 1 |
| 10N26155C8 | 1.281.O | 10N26155C8_0_2482 | Na(+)-translocating NADH-quinone reductase subunit B (EC 1.6.5.-) | 1 |
| 10N26155C8 | 1.281.O | 10N26155C8_0_2480 | Na(+)-translocating NADH-quinone reductase subunit D (EC 1.6.5.-) | 1 |

**Table S3.**
Receptor identification by sequencing of spontaneous resistant mutants.

| Host | Phage | Replicate | Contig_Position | Type | Length | Gene | AA change | Annotation |
|---|---|---|---|---|---|---|---|---|
| 10N26155C8 | 1.119.O | 1 | 2_171942 | Deletion | 1 | ORF_2_160 | ORF_2_160:p.Lys7fs | General secretion pathway protein G |
| 10N26155C8 | 1.119.O | 2 | 2_173977 | SNV | 1 | ORF_2_162 | ORF_2_162:p.Lys267* | General secretion pathway protein E |
| 10N26155C8 | 1.119.O | 3 | 2_1671-167305 | Deletion via CA | 109 | ORF_2_154 | NA | General secretion pathway protein M |
| 10N26155C8 | 1.119.O | 4 | 2_167254 | SNV | 1 | ORF_2_154 | ORF_2_154:p.Trp63* | General secretion pathway protein M |
| 10N26155C8 | 1.119.O | 5 | 2_173221 | Deletion | 1 | ORF_2_161 | ORF_2_161:p.Gly18fs | General secretion pathway protein F |
| 10N26155C8 | 1.119.O | 6 | 2_1719-172018 | Deletion via CA | 83 | ORF_2_160 | NA | General secretion pathway protein G |
| 10N26155C8 | 1.119.O | 6 | 2_166050-166174 | Deletion via CA | 125 | ORF_2_153 | NA | General secretion pathway protein N |
| 10N26155C8 | 1.281.O | 1 | 2_2804974 | SNV | 1 | ORF_0_2483 | ORF_0_2483:p.Ser302* | Na(+)-translocating NADH-quinone reductase subunit A (EC 1.6.5.-) |
| 10N26155C8 | 1.281.O | 2 | 2_68299 | Insertion | 1 | ORF_2_61 | ORF_2_61:p.Tyr187fs | Unannotated |
| 10N26155C8 | 1.281.O | 3 | 2_72111-72115 | Deletion via CA | 5 | NA | NA | intergenic, upstream of 10N26155C8_2_65 (Unannotated) |
| 10N26155C8 | 1.281.O | 3 | 2_72128-72134 | Deletion via CA | 7 | NA | NA | intergenic, upstream of 10N26155C8_2_65 (Unannotated) |
| 10N26155C8 | 1.281.O | 4 | 2_68299 | Insertion | 1 | ORF_2_61 | ORF_2_61:p.Tyr187fs | Unannotated |
| 10N26155C8 | 1.281.O | 5 | 2_108233 | SNV | 1 | ORF_2_96 | ORF_2_96:p.Gln332* | dTDP-glucose 4,6-dehydratase (EC 4.2.1.46) |
| 10N26155C8 | 1.281.O | 6 | 2_68845-68906 | Deletion via CA | 62 | ORF_2_61&62 | NA | end of 10N26155C8_2_61 and beginning of 10N26155C8_2_62 |
| 10N26155C8 | 1.281.O | 7 | 2_107805 | SNV | 1 | ORF_2_95 | ORF_2_95:p.Asn109Tyr | Glucose-1-phosphate thymidylyltransferase (EC 2.7.7.24) |

**Table S4.**

Predicted motifs of RM systems on PDEs and methylation fraction in genome.

| | | |
|---|---|---|
| RM PDE | PDE 1 | |
| Predicted Motif | TCA*BN(4)RTRTC | |
| | Host/Phage | Fraction Methylated |
| | 10N26155C8 | 599/600 |
| | 1.119.O | 1/1 |
| | 1.127.O | 1/1 |
| | 1.143.O | 1/1 |
| | 1.231.O | 1/1 |
| | 10N28654F7 | 0/599 |
| | 1.283.A | 0/7 |
| | 1.281.O | 0/8 |
| | 1.196.O | 0/8 |
| RM PDE | PDE 4 | |
| Predicted Motif | CCA*GN(6)TAA | |
| | Host/Phage | Fraction Methylated |
| | 10N26155C8 | 0/646 |
| | 1.119.O | 0/3 |
| | 1.127.O | 0/4 |
| | 1.143.O | 0/3 |
| | 1.231.O | 0/3 |
| | 10N28654F7 | 635/638 |
| | 1.283.A | 5/5 |
| | 1.281.O | 4/4 |
| | 1.196.O | 4/4 |
| RM PDE | PDE 5 | |
| Predicted Motif | GA*GN(6)GGC | |
| | Host/Phage | Fraction Methylated |
| | 10N26155C8 | 0/1795 |
| | 1.119.O | 0/17 |
| | 1.127.O | 0/15 |
| | 1.143.O | 0/17 |
| | 1.231.O | 0/17 |
| | 10N28654F7 | 1770/1779 |
| | 1.283.A | 8/8 |
| | 1.281.O | 10/10 |
| | 1.196.O | 10/10 |

**Table S5.**

Strains and plasmids used in transposon mutagenesis and gene deletions.

| Strain or plasmid | Description | Reference |
|---|---|---|
| **Strains** | | |
| ***E. coli*** | | |
| β3914 | ($F^-$) RP4-2-Tc::Mu *ΔdapA*::(erm-pir) *gyrA462 zei-298*::Tn10 ($Km^R$ $Erm^R$ $Tc^R$) | (*63*) |
| Π3813 | *lacI^q thi-1 supE44 endA1 recA1 hsdR17 gyrA462 zei298::Tn10 ΔthyA::(erm-pir-116)* ($Tc^R$ $Erm^R$) | (*63*) |
| MFD*pir* | *E. coli* MG1655 RP4-2-Tc::[ΔMu1::*aac(3)IV*-Δ*aphA*-Δ*nic35*-ΔMu2*::zeo*] Δ*dapA*::(erm-pir) Δ*recA* ($Apra^R$ $Zeo^R$ $Erm^R$) | (*61*) |
| ***V. lentus*** | | |
| 10N.261.55.C8 | Representative "orange" strain (C8-WT) | This study |
| ΔPDE1 | C8-WT with ΔPDE1: in frame partial deletion of PDE1 (31909/34140 bp) | This study |
| ΔPDE2 | C8-WT with ΔPDE2: in frame partial deletion of PDE2 (12186/20725 bp) | This study |
| ΔPDE3 | C8-WT with ΔPDE3: in frame partial deletion of PDE3 (25697/37369 bp) | This study |
| ΔΔPDE12 | C8-WT with ΔPDE1 and ΔPDE2 | This study |
| ΔΔPDE13 | C8-WT with ΔPDE1 and ΔPDE3 | This study |
| ΔΔPDE23 | C8-WT with ΔPDE2 and ΔPDE3 | This study |
| ΔΔΔPDE123 | C8-WT with ΔPDE1, ΔPDE2 and ΔPDE3 | This study |
| **Plasmids** | | |
| pSC189-Cm | *oriT RP4* Π-dependent *oriV* R6K *mariner*-based transposon TnSC189 Δ*kan::cat* ($Cm^R$ $Ap^R$) | (*61*) |
| pSW7848T | *oriV R6K ; oriT RP4; araC-$P_{BAD}$ccdB $Cm^R$* | (*75*) |
| pSWδR-1 | pSW7848T::ΔPDE1 | This study |
| pSWδR-2 | pSW7848T::ΔPDE2 | This study |
| pSWδR-2 | pSW7848T::ΔPDE3 | This study |

**Table S6.**
Primers used in transposon mutagenesis and gene deletions.

| Primer | Sequence 5´-3´ | Reference |
|---|---|---|
| SS9arb2 | GACCACGAGACGCCACACTNNNNNNNNNNACTAG | (*65*) |
| Mar4 | TAGGGTTGAGTGTTGTTCCAGTT | (*66*) |
| Mar4_int2 | GTCATCGTCATCCTTGTAATCG | This study |
| Arb3 | GACCACGAGACGCCACACT | (*65*) |
| ΔPDE1/F1 | GTCGACGGTATCGATAAGCTTGATATCGAATTCCTGCATCATGGCTTGG GTCACTCG | This study |
| ΔPDE1/R1 | GAAACTGGGTGCAAATGTCGTACAGTCTGGTGGGCCTGAG | This study |
| ΔPDE1/F2 | CTCAGGCCCACCAGACTGTACGACATTTGCACCCAGTTTC | This study |
| ΔPDE1/R2 | CCGTCAAGTTGTCATAATTGGTAACGAATCAGACAATTTTGTACCCTAGC GAACATTCTG | This study |
| ΔPDE1/F | GCCTACAGGTTGCTTTCGTC | This study |
| ΔPDE1/R | CAGCGCGTATTCTCTCGTTG | This study |
| ΔPDE2/F1 | TAAGCTTGATATCGAATTCCTGCAGGTTGCCATCATTCTATTCGG | This study |
| ΔPDE2/R1 | TGTTAAGGAAGTGGCAAAGTGAATGCACCAAGACTCACCACGAAG | This study |
| ΔPDE2/F2 | AAACCACTTCGTGGTGAGTCTTGGTGCATTCACTTTGCCACTTCC | This study |
| ΔPDE2/R2 | AATTGGTAACGAATCAGACAATTTTGTGAGAAGTACGGTGTTTGG | This study |
| ΔPDE2/F | TCGCTGAGGTTTGCTCTAC | This study |
| ΔPDE2/R | ATTACGATGAAGCTCAAAGCC | This study |
| ΔPDE3/F1 | GCTTGATATCGAATTCCTGCAATTGCTAACCTACTGCCTTAC | This study |
| ΔPDE3/R1 | GGAAGTGGCAAAGTGAATGCTGGAAACTCACTCACTCACTC | This study |
| ΔPDE3/F2 | GAGTGAGTGAGTGAGTTTCCAGCATTCACTTTGCCACTTCC | This study |
| ΔPDE3/R2 | CATAATTGGTAACGAATCAGACAATTGATGCTTATCGTGCGGTAAATG | This study |
| ΔPDE3/F | GCGTAATGTCAGTTTGATTTCGATG | This study |
| ΔPDE3/R | CAAGATCACTATGCAGGAACAGG | This study |
| pSW_ F | AATTGTCTGATTCGTTACCAATTATG | This study |
| pSW_ R | TGCAGGAATTCGATATCAAGC | This study |

**Data S1.**
Detailed gene annotations for key phage defense elements in Figure 1B.

**Data S2A.**
FASTA file of putative phage defense regions presented in Figure S8.

**Data S2B.**
Map for Data S2A in CSV format.

**Data S3B.**
FASTA file of putative phage defense regions presented in Figure S10.

**Data S3B.**
Map for Data S3A in CSV format.

**Data S4B.**
FASTA file of putative phage defense regions presented in Figure S11.

**Data S4B.**
Map for Data S4A in CSV format.
